# Cortico-amygdalar connectivity and externalizing/internalizing behavior in children with neurodevelopmental disorders

**DOI:** 10.1101/2021.02.01.429143

**Authors:** Hajer Nakua, Colin Hawco, Natalie J. Forde, Grace R. Jacobs, Michael Joseph, Aristotle Voineskos, Anne L. Wheeler, Meng-Chuan Lai, Peter Szatmari, Elizabeth Kelley, Xudong Liu, Stelios Georgiades, Rob Nicolson, Russell Schachar, Jennifer Crosbie, Evdokia Anagnostou, Jason P. Lerch, Paul D. Arnold, Stephanie H. Ameis

## Abstract

**Background:** Externalizing and internalizing behaviors are common and contribute to impairment in children with neurodevelopmental disorders (NDDs). Associations between externalizing or internalizing behaviors and cortico-amygdalar connectivity have been found in children with and without clinically significant internalizing/externalizing behaviors. This study examined whether such associations are present across children with different NDDs.

**Methods:** Multi-modal neuroimaging and behavioral data from the Province of Ontario Neurodevelopmental Disorders (POND) Network were used. POND participants aged 6-18 years with a primary diagnosis of autism spectrum disorder (ASD), attention-deficit/hyperactivity disorder (ADHD) or obsessive-compulsive disorder (OCD), as well as typically developing children (TDC) with T1-weighted, resting-state fMRI or diffusion weighted imaging and parent-report Child Behavioral Checklist (CBCL) data available, were analyzed (n range=157-346). Associations between externalizing or internalizing behavior and cortico-amygdalar structural and functional connectivity indices were examined using linear regressions, controlling for age, gender, and image-modality specific covariates. Behavior-by-diagnosis interaction effects were also examined.

**Results:** No significant linear associations (or diagnosis-by-behavior interaction effects) were found between CBCL-measured externalizing or internalizing behaviors and any of the connectivity indices examined. Post-hoc bootstrapping analyses indicated stability and reliability of these null results.

**Conclusions:** The current study provides evidence in favour of the absence of a shared linear relationship between internalizing or externalizing behaviors and cortico-amygdalar connectivity properties across a transdiagnostic sample of children with various NDDs and TDC. Detecting shared brain-behavior relationships in children with NDDs may benefit from the use of different methodological approaches, including incorporation of multi-dimensional behavioral data (i.e. behavioral assessments, neurocognitive tasks, task-based fMRI) or clustering approaches to delineate whether subgroups of individuals with different brain-behavior profiles are present within heterogeneous cross-disorder samples.

## INTRODUCTION

Autism spectrum disorder (ASD), attention-deficit/hyperactivity disorder (ADHD), and pediatric obsessive-compulsive disorder (OCD) are neurodevelopmental disorders (NDDs) with high rates of clinical co-occurrence (Abramovitch et al., 2015; Jang et al., 2013; Lai et al., 2019; Lewin et al., 2011; Masi et al., 2006) in addition to significant overlap in clinical (Lawson et al., 2015; Mito et al., 2014), behavioral (Anholt et al., 2010; Havdahl et al., 2016), cognitive (Antshel et al., 2013; Van Der Meer et al., 2012), genetic (Lionel et al., 2014, 2011), and brain features (Ameis et al., 2016; Kern et al., 2015). This overlap has motivated recent transdiagnostic research examining the shared and distinct biological and behavioral features across these disorders (Ameis et al., 2016; Carlisi et al., 2017; Kushki et al., 2019). Externalizing (e.g., aggression, rule-breaking) and internalizing (e.g., withdrawal, anxiety, depression, somatic) behaviors manifest across children and youth to varying degrees (Bradley et al., 2004; Dwyer et al., 2006; Ghandour et al., 2019; Jacob et al., 2014). Children and youth with NDDs are more likely to exhibit clinically significant behaviors in either domain (Alvarenga et al., 2016; Bauminger et al., 2010; Jacob et al., 2014), contributing to increased functional impairment (e.g., at school and home) (Arim et al., 2015; Mazurek et al., 2013) and poorer response to interventions (Hill et al., 2014; Torp et al., 2015).

Internalizing and externalizing behaviors have been linked to a cortico-amygdalar network comprised of frontal cortical regions implicated in decision-making, behavioral regulation (Rushworth et al., 2011), and emotional regulation (Albaugh et al., 2016; Ducharme et al., 2011) which provide top-down modulation of amygdalar activity (Etkin et al., 2006; Hariri et al., 2003). This cortico-amygdalar network is connected through two main white matter tracts: the uncinate fasciculus (UF) and the cingulum bundle (CB) (Catani et al., 2013). In typically developing children (TDC), increased internalizing behavior has been associated with altered structural covariance between the prefrontal cortex and amygdala (Vijayakumar et al., 2017), decreased fractional anisotropy (FA) of the UF and CB (Albaugh et al., 2016; Mohamed Ali et al., 2019), and increased functional connectivity between the ventromedial prefrontal cortex and amygdala (Qin et al., 2014). Also in TDC, increased externalizing behavior has been associated with altered cortico-amygdalar structural covariance (Ameis et al., 2014), decreased FA of the UF (Andre et al., 2019), and altered functional connectivity between amygdala and frontal cortical regions (Aghajani et al., 2017, 2016; Saxbe et al., 2018). These studies suggest that cortico-amygdalar connectivity may be associated with both externalizing and internalizing behaviors, which often co-occur (Korhonen et al., 2014; Reef et al., 2011). Cortico-amygdalar network alterations have also been found in studies of children with primary internalizing (e.g., major depressive disorder) or externalizing (e.g., oppositional defiant disorder) disorders (Castellanos-Ryan et al., 2014; Luking et al., 2011; Noordermeer et al., 2016; Paulesu et al., 2010; Stoycos et al., 2017) compared to TDC. Shared continuous associations between task-based fMRI and behavioral measures (parent report and in-scanner assessments) have also been found across children with different psychiatric diagnoses (i.e., disruptive behavior disorders, anxiety disorders, or ADHD) (Ibrahim et al., 2019; Stoddard et al., 2017). Taken together, these studies suggest that cortico-amygdalar connectivity properties may relate to internalizing or externalizing behaviors along a continuum and cutting across TDC and children and youth with different mental health diagnoses.

As yet, we know of no study that has investigated whether cortico-amygdala network properties relate to internalizing or externalizing behaviors across children with different NDDs. The present study investigated linear associations between externalizing or internalizing behaviors and indices of cortico-amygdalar network connectivity (i.e., structural covariance, resting-state functional connectivity, and white matter connectivity) in a large sample, including TDC and children and youth with primary diagnoses of ASD, OCD, or ADHD. We hypothesized that greater externalizing or internalizing behaviors would be associated with reduced cortico-amygdalar structural and functional connectivity indices across our transdiagnostic sample.

## METHODS

### Participants

Participants included in the current study participated in the Province of Ontario Neurodevelopmental Disorders (POND) Network; recruitment was carried out at different sites across the province of Ontario, Canada, including the Hospital for Sick Children (SickKids), Holland Bloorview Kids Rehabilitation Hospital, Lawson Health Research Institute, McMaster University and Queen’s University between June 2012 and January 2020. Children and youth were eligible to participate in POND if they had a primary clinical diagnosis of ASD, ADHD or OCD, sufficient English language comprehension to complete the behavioral assessments, and no contraindications for MRI (e.g., metal implants). The Parent Interview for Child Symptoms (Ickowicz et al., 2006) was used to confirm ADHD diagnosis, the Schedule for Affective Disorders-Children’s Version (Kiddie-SADS) and Children’s Yale-Brown Obsessive-Compulsive Scale (Scahill et al., 1997) for OCD, and the Autism Diagnostic Observation Schedule-2 (Lord et al., 2000) and the Autism Diagnostic Interview-Revised (Disorders et al., 1994) for ASD. TDC participants were recruited through flyers posted at each recruitment site as well as through word-of-mouth. The exclusion criteria for TDC included: history of premature birth (<35 weeks), presence of an NDD, first-degree relative with an NDD, psychiatric or neurologic diagnosis, confirmed via parental screening. Age-appropriate Wechsler scales were used to estimate full-scale IQ (Littell, 1960). Participating institutions received approval for this study from their respective research ethics boards. Primary caregivers and participants provided either written informed consent or assent after a complete description of the study was provided. As of January 2020, MRI and behavioral data were available for 611 children and youth with ASD, ADHD, OCD, or TDC (n=286 ASD; n=159 ADHD; n=68 OCD; n=98 TDC) who completed MRI scanning at SickKids (Toronto, Canada). The present study analyzed data from a subset of these 611 POND participants who met all of the following criteria: (i) they had successfully completed a T1-weighted, resting-state, or single-shell DWI scan, (ii) were between the ages of 6 and 18 years at time of brain scan, and (iii) had Child Behavior Checklist (CBCL) data available that was collected within 12 months of their MRI scan (see *Figure 1*).

**Figure 1.**
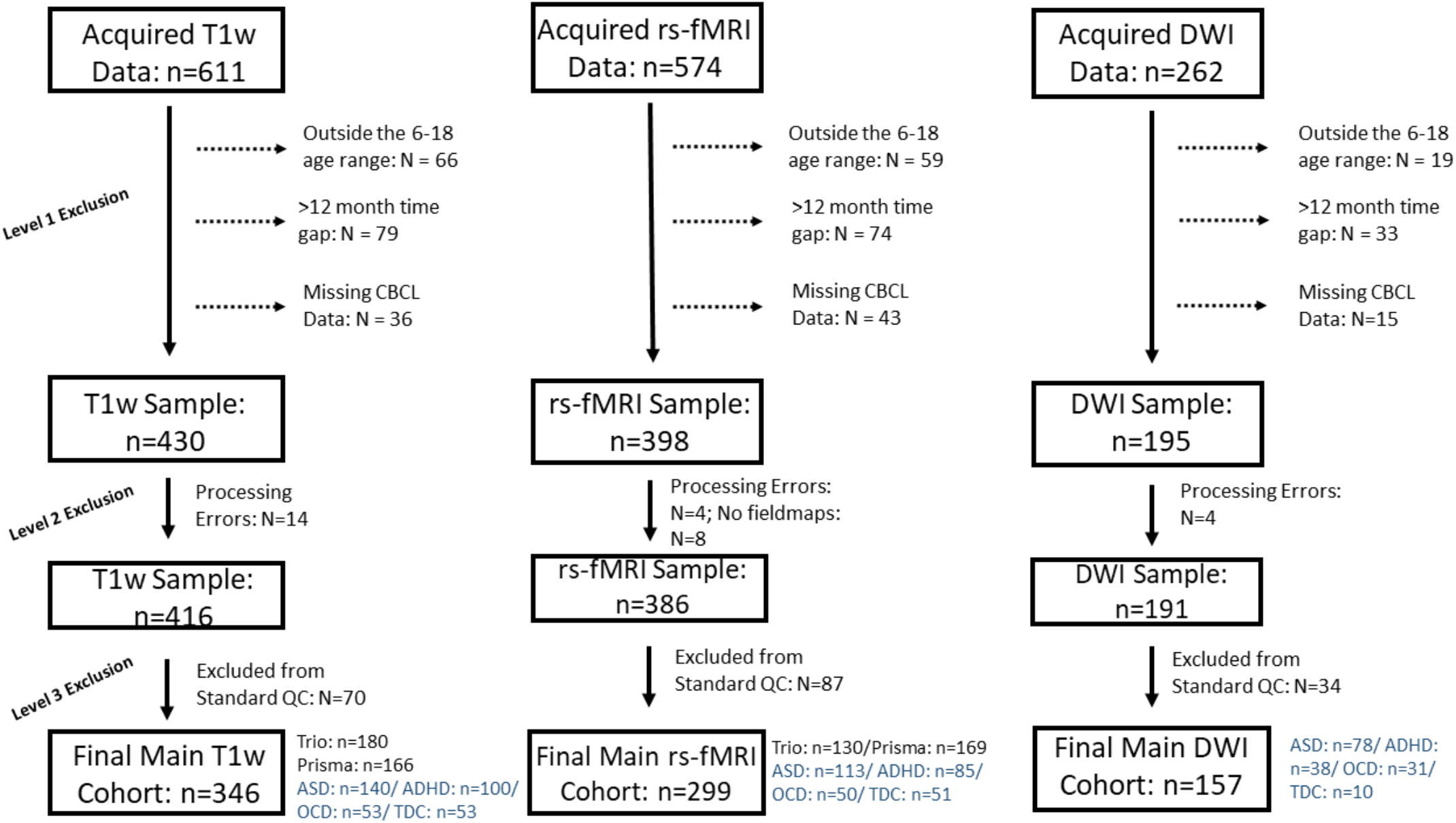
Diagrams presenting the overall POND imaging samples which includes children with autism spectrum disorder (ASD), attention-deficit/hyperactivity disorder (ADHD), obsessive compulsive disorder (OCD) and typically developing children (TDC) scanned at the Hospital for Sick Children as of January 2020. Imaging data from T1-weighted (T1w), resting state fMRI (rsfMRI) and diffusion weighted imaging (DWI) sequences are presented. The reasons for exclusion presented for level 1: participants being outside the 6-18 age range at time of scan, a greater than 12 month time gap between scan and CBCL administration, and missing CBCL data; level 2: persistent processing errors at any point within the processing pipeline (e.g. errors in the fMRIprep pipeline); level 3: exclusion based on quality control (QC; details presented in the paper and supplement). The numbers for the final analysed sample for each imaging modality are presented. For the T1w and rs-fMRI samples, participants were scanned on a 3T Siemens Tim Trio scanner prior to June 2016 when the scanner was upgraded to the PrismaFIT. For rs-fMRI acquisitions, participants scanned on the Tim Trio selected a movie to watch and participants scanned on the PrismaFIT viewed a naturalistic film (inscapes). The study includes only single-shell DWI acquisitions (n=262) completed on the Tim Trio scanner.

### Measurement of Externalizing and Internalizing Behaviors

Externalizing and internalizing behavioral scores were measured using the parent-report CBCL (ages 6-18), a standardized, well-established instrument (Achenbach and Ruffle, 2000) that has been widely used for brain-behavior analyses in pediatric samples (Albaugh et al., 2016; Ameis et al., 2014; Ducharme et al., 2014, 2011; Ibrahim et al., 2019). The CBCL provides continuous measures of externalizing (calculated by combining rule-breaking and aggressive CBCL subscales) and internalizing behavior (calculated by combining withdrawn, anxious/depressed and somatic CBCL subscales), with a domain specific t-score (standardized by age and gender) >70 indicating clinically significant symptoms.

### MRI Protocol

Participants were scanned on a 3T Siemens Tim Trio at SickKids that was upgraded to the Siemens PrismaFIT in June 2016. All T1-weighted brain imaging consisted of a 5-minute scan using an MPRAGE sequence with grappa parallelization (Tim Trio: (1×1×1)mm^3^, TR=2,300ms, TE=2.96ms, TI=900ms, Flip Angle=9°, FOV=224×224mm^2^, 240 Slices, GRAPPA=2, 12-channel head coil; PrismaFIT: (0.8×0.8×0.8)mm^3^, TR=1,870ms, TE=3.14ms, TI=945ms, Flip Angle=9°, FOV=222×222mm^2^, 240 Slices, GRAPPA=2, 20-channel head coil). Resting state functional MRI (rs-fMRI) data consisted of a ∼5-minute scan (Tim Trio: (3.5×3.5×3.5)mm^3^, TR=2340ms, TE=30ms, Flip Angle=70°, FOV=256×240mm^2^, 120 volumes; PrismaFIT: (3×3×3)mm^3^, TR=1500ms, TE=30ms, Flip Angle=70° FOV=256×240mm^2^, 200 volumes). During rs-fMRI scanning, participants either viewed a movie of their choice, if the scan occurred pre-upgrade on Tim Trio, or a naturalistic movie paradigm (Vanderwal et al., 2015), if the scan occurred post-upgrade on PrismaFIT.

Single-shell DWI scans were acquired as 3 consecutive sequences with 19, 20 or 21 gradient directions (for a total of 60 directions) and 3 B0’s per acquisition sequence. Scan parameters were as follows: ((2×2×2)mm3, TR=3800ms, TE=73ms, Flip Angle=90° FOV=244×244mm2, 69 volumes, B0=1000). Multi-shell DWI data were acquired post scanner upgrade to PrismaFIT. Multi-shell data were not analyzed in the current study due to the challenges of harmonization across different DWI scan acquisition protocols and concerns regarding measurement variability given substantial differences in the pre-to-post hardware upgrade sequence design (Tax et al., 2019).

### Image Pre-processing

Prior to pre-processing, the scans of participants who had multiple image acquisitions underwent quality assessment and the higher quality scan was pre-processed. Visual examination of the raw brain scan was used to assess the quality of T1-weighted and DWI acquisitions. Quality metric comparisons (e.g., mean framewise displacement [FD]) from the MRIQC pipeline (Esteban et al., 2017) was used to assess rs-fMRI acquisitions.

### Structural MRI

T1-weighted brain images were pre-processed using the fMRIprep pipeline (Esteban et al., 2019) which runs FreeSurfer and performs intensity non-uniformity correction, skull stripping, calculates spatial normalization based on an MNI template, tissue segmentation and surface reconstruction. Images were also run through the MRIQC pipeline (Esteban et al., 2017) to extract quality metrics used in the quality control (QC) procedure. Left and right amygdala volumes from each participant were extracted using the amygdala region-of-interest (ROI) defined by the Desikan-Killiany Atlas (Desikan et al., 2006). The ciftify pipeline ((Dickie et al., 2019); https://github.com/edickie/ciftify) was used to transform the images from the FreeSurfer format to the Connectivity Informatics Technology Initiative (CIFTI) format. From there, the 40,962 vertices in each hemisphere were extracted based on FreeSurfer’s white and pial surfaces. This pipeline registered cortical surfaces to an average surface to establish correspondence between participants. Cortical thickness values at each vertex were smoothed with a Gaussian kernel of 12mm full width half maximum (FWHM).

### Resting state fMRI

The rs-fMRI acquisitions were pre-processed through fMRIPrep (Esteban et al., 2019). Within fMRIPrep, the data was slice timed and motion corrected. Distortion correction was performed using field maps; the functional image was co-registered to the corresponding T1-weighted image using FreeSurfer with boundary-based registration with 9 degrees of freedom. Nonlinear transformation to the MNI152 template was calculated via FSL’s FNIRT (based on the T1-weighted image) and applied to the functional data. These data were then transformed onto the cortical surface and converted to the CIFTI format (Dickie et al., 2019). The first three TRs were dropped, and voxel time series underwent mean-based intensity normalization, linear and quadratic detrending, temporal bandpass filtering (0.009-0.08 Hz), and confound regression for 6 head motion parameters, white matter signal, CSF signal and global signal plus their lags, their squares, and the squares of their lags (i.e. a 24HMP + 8 Phys + 4 GSR confounds) (Ciric et al., 2018). Global signal regression was employed as it has been shown to reduce sources of noise and reduce correlations between mean FD and functional connectivity (Parkes et al., 2018). Spatial smoothing was then performed on the cortical surface data (FWHM=8mm).

### Diffusion Weighted Imaging

DWI scans from the three separate runs were concatenated. Diffusion data were denoised using random field theory and upsampled to a (1×1×1) mm^3^ voxel size using the *MRtrix3* dwidenoise and *mrresize* commands, respectively (Veraart et al., 2016). Using fieldmaps, images were corrected for motion artefacts accounting for field inhomogeneities and eddy current induced artefacts using FSL’s (Smith et al., 2004) eddy function (Andersson et al., 2016; Andersson and Sotiropoulos, 2016). Deterministic tractography was used to delineate the UF and CB via the Slicer dMRI software (https://github.com/SlicerDMRI). The software registered the tracts for all participants using a dataset-specific atlas based on a representative subset (n=21) from the current sample (selected based on age, gender and diagnosis) (Fedorov et al., 2012). Within the Slicer software, fiber clusters were manually appended to create the white matter tracts of interest (CB and UF). The atlas was registered to all participants’ DWI acquisitions and FA and mean diffusivity (MD) metrics were extracted.

### Quality Control (QC)

To reduce potential bias of image artefacts, a rigorous *a priori* QC procedure was applied for all imaging modalities (supplementary section 1; *Figure S1*). T1-weighted images were assessed for motion artefacts using a visual QC (HTML visual outputs from the fMRIPrep pipeline) in addition to quantitative QC (MRIQC-derived quality metrics). For the rs-fMRI sequence, participants that did not complete the ∼5-min scan were excluded based on prior research indicating this time duration is required for stable estimations of correlation strengths (Van Dijk et al., 2010). Quality of rs-fMRI acquisitions were assessed based on mean FD and excluded based on in-scanner motion at mean FD>0.5mm given that clinical sample have greater in scanner head motion (Pardoe et al. 2016). DWI acquisitions were assessed for slice dropouts, poor V1 directions and residuals using an in-house standardized pipeline in addition to quantitative quality metrics. See *Figure S1* and supplement for QC procedure details.

### Data and Code Availability

See https://github.com/hajernakua/cortico-amygdalar2019 for the code used in our analyses (pre-processing and analysis scripts) and QC procedures. The POND Network has made a commitment to release the POND data. Data release is controlled and managed by the Ontario Brain Institute (OBI). The OBI POND data will be released via the Brain-CODE portal. For further details please see https://www.braincode.ca/.

## Statistical Analysis

### Brain-behavior relations

Separate linear regression models were fit to examine the presence of an association between externalizing or internalizing behavioral scores and cortico-amygdalar connectivity metrics, including: structural covariance, seed based rs-fMRI as well as FA and MD of the UF and CB (see below and supplementary for details). Covariates for the primary regression models included age, gender, and scanner (if acquisitions from a modality included a scanner upgrade). Across image modalities, if the primary regression models were significant, subsequent models were planned to sequentially fit the following covariates: (i) the alternate broad-band CBCL score (e.g., internalizing behavior as a covariate when externalizing behavior is the predictor variable) to account for shared variability (Zald & Lahey, 2017), (ii) mean FD for functional connectivity models (Power et al., 2012; Satterthwaite et al., 2012) or an estimate of overall noise for white matter connectivity models (Anderson, 2001) (*see details in S3.2.1*), and (iii) medication status (taking medication, not taking medication, unknown).

### Brain/behavior age relations

Prior to fitting the brain-behavior structural covariance, functional connectivity and white matter connectivity models, linear regression models were fit between age, age-squared, and brain and behavior indices. The better fitting age term was included as a covariate in the main analyses (*Table S2; supplementary section 5.1*). If the better fitting age term was quadratic, then linear and quadratic age terms were included in the model.

### Cortico-amygdalar structural covariance

Using a partial regression, an interaction term (independent variable) between internalizing or externalizing behavior scores and left or right amygdala volume (e.g., externalizing behavior*left amygdala volume) was regressed onto each cortical vertex (with thickness at each vertex as the dependent variable) controlling for age, parent reported gender (male, female), intracranial volume (Buckner et al., 2004; Raz et al., 2004) and scanner (i.e., Tim Trio pre-upgrade or PrismaFIT post-upgrade). Analyses were carried out using FSL’s Permutation Analysis of Linear Models (PALM) package. Clusters of vertex-wise significance were determined using 2000 permutations with the threshold free cluster enhancement (TFCE) approach (Smith and Nichols, 1996). Considering the high number of cortical vertices and consequent linear models, group results were thresholded at p<0.05 FDR-corrected from the number of vertices in each hemisphere, and further corrected for separate runs of PALM for each hemisphere (critical level *a*=0.025). Eight models were fit with cortical thickness at each vertex as the dependent variable to account for the different combinations between behavioral scores and amygdala volume and the behavior-by-diagnosis terms. Similar partial regression models were used for analyses examining rs-fMRI and DWI metrics as dependent variables. See below an example of one of the linear regression models examining associations between left cortico-amygdalar structural covariance and externalizing behavior.

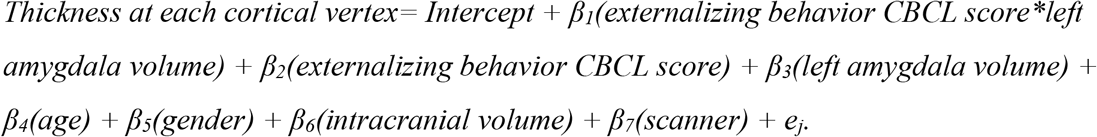

### Resting-state functional connectivity

Seed-based functional connectivity analyses were performed using the left and right amygdala as seed ROIs. Mean time series for each amygdala ROI were correlated with the time series of each cortical vertex using the *ciftify_seed_corr* function from the ciftify pipeline. Externalizing or internalizing behavior was regressed onto the functional connectivity between the amygdala ROI and each cortical vertex, controlling for age, gender, and scanner. PALM with TFCE was used to control for multiple comparisons across cortical vertices.

### White matter connectivity

Using R (version 3.5.0), internalizing and externalizing behavioral scores were regressed onto left or right UF and CB for FA and MD metrics, controlling for age and gender. An FDR correction was applied to the primary analyses examining associations between behavior and left or right CB and UF diffusion metrics.

### Interaction Effects

Behavior-by-diagnosis interaction terms were fit in separate models to examine whether brain-behavior relationships were influenced by diagnostic status.

### Planned Subsample Analysis

Given the potential for considerable behavioral and brain change over time in a developing sample (Bos et al., 2018), a sensitivity analysis was conducted in a subset of participants whose brain scan was obtained within one month of completion of their CBCL data (see *Figure S2* in supplemental materials for subsample details).

### Post-hoc bootstrap resampling analysis

The linear brain-behavior models examined in the current study were not significant (see results section). The lack of replicability of neuroimaging research findings (Ioannidis, 2018; Simmons et al., 2011; He et al. 2020) has led to calls for increased efforts to assess reliability of reported results (Button et al. 2013). Null results have an additional challenge of interpretability as a null effect does not necessarily indicate the absence of an effect. Here, we used bootstrap resampling to assess the stability and reliability of the null effect for the brain-behavior models which address the main aims of the current study (i.e., models that examined the main effect of externalizing or internalizing behavior across cortico-amygdalar connectivity indices in the current sample). Using a case-resampling bootstrap (Monte Carlo) approach, 1000 iterations of each data (i.e., design) matrix were generated and used to perform repeated linear regressions for each generated sample. Each iteration of the data matrix randomly selected participants with replacement until the total sample size was reached (e.g., n=346 for the main structural covariance models). Stability of the models were assessed using the bootstrap resampled standard errors of the regression coefficients (McIntosh & Lobaugh, 2004; Efron & Tibshirani, 1986). The reliability of assessed models was assessed by examining the distributions (i.e., standard deviations) of the resampled model parameter estimates (i.e., regression coefficients, t-statistics, and effect sizes; Himberg et al. 2004). Stable and reliable models feature near-zero standard errors and low standard deviations of parameter estimates (McIntosh & Lobaugh, 2004; Efron & Tibshirani, 1986). For the structural covariance and functional connectivity models, bootstrap resampling was conducted in PALM and parameter estimates were calculated at each vertex. For the white matter connectivity models, each model was analyzed in RStudio and the standard errors of each model regression coefficients were calculated (see supplementary section 7 for more details about the bootstrap resampling analysis).

## RESULTS

### Participant Information

Characteristics of the analyzed sample are shown in *Table 1*. Following removal of participants outside the 6-18 age range, those with a time window greater than 12 months between acquired scan and behavioral assessment date and those who failed QC, a total of 346 participants were included for structural covariance, 299 participants for resting-state functional connectivity and 157 participants for white matter connectivity analyses (see *Figure 1* for consort diagram). Across all image modalities, there were no significant differences in externalizing and internalizing behavior scores between the total POND SickKids sample scanned by January 2020 (n=611) and the subsample analyzed in the current study, nor were there any differences in age, gender, or diagnostic composition (see supplementary section 4 for further details). A significant positive correlation between age and internalizing behaviors was found across the analyzed samples (i.e. T1w, rsfMRI, and DWI samples; see supplementary section 4). A significant negative correlation between age and externalizing behaviors was found in the T1w sample (r=-0.11, p=0.044). There were no significant differences in age (F=1.44, p=0.24), internalizing (F=0.03, p=0.87) or externalizing (F=0.41, p=0.66) behaviors between participants included in T1w, rs-fMRI, and DWI samples. In the T1 and rs-fMRI samples, there was a significant difference in diagnostic composition across the scanner upgrade (X^2^=32.48, p<0.001), as more TDC participants were scanned post upgrade.

**Table 1.**
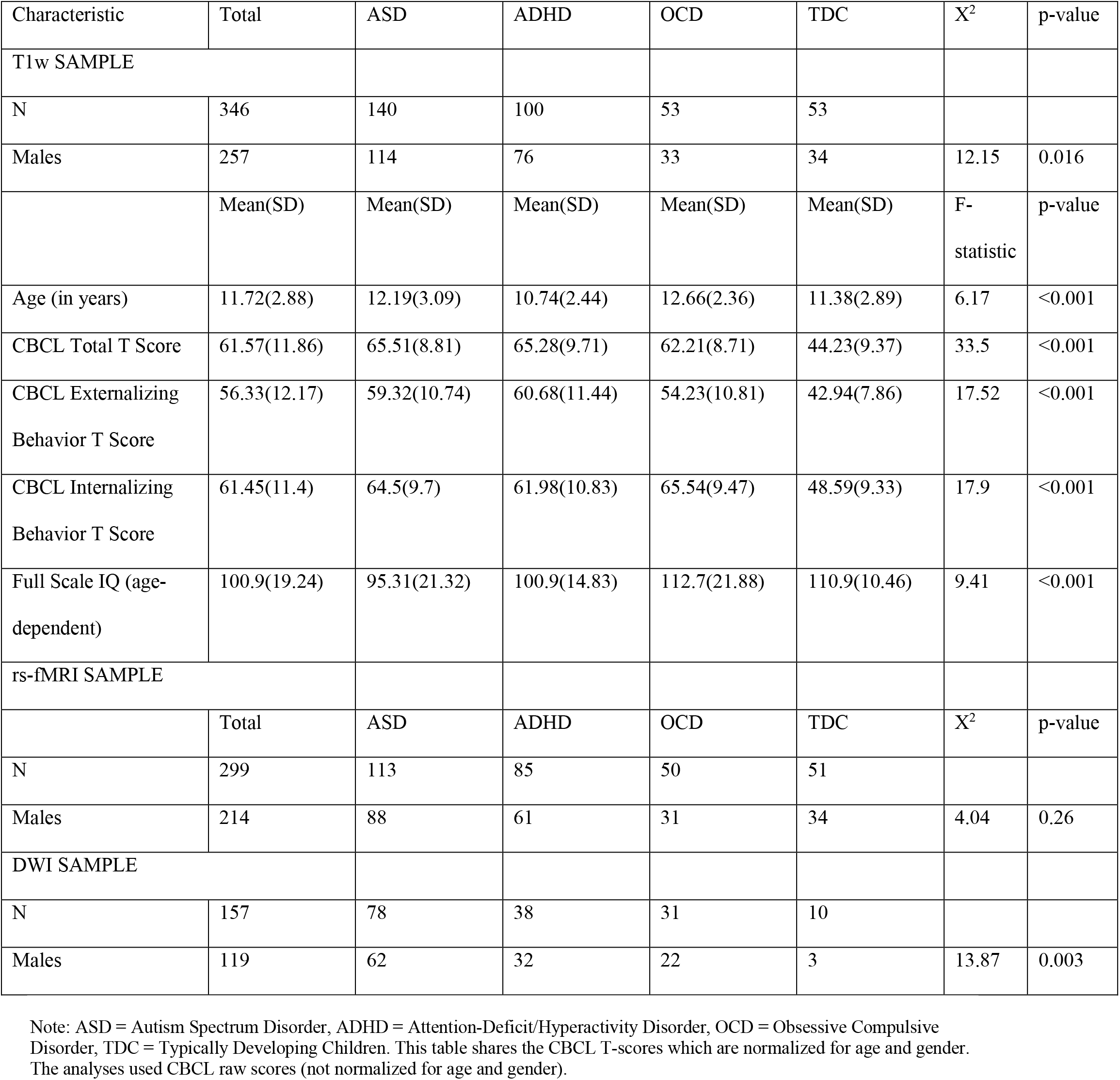
Demographic Characteristics of the data analyzed from the POND dataset

| Characteristic | Total | ASD | ADHD | OCD | TDC | X<sup>2</sup> | p-value |
| --- | --- | --- | --- | --- | --- | --- | --- |
| T1w SAMPLE |  |  |  |  |  |  |  |
| N | 346 | 140 | 100 | 53 | 53 |  |  |
| Males | 257 | 114 | 76 | 33 | 34 | 12.15 | 0.016 |
|  | Mean(SD) | Mean(SD) | Mean(SD) | Mean(SD) | Mean(SD) | F-statistic | p-value |
| Age (in years) | 11.72(2.88) | 12.19(3.09) | 10.74(2.44) | 12.66(2.36) | 11.38(2.89) | 6.17 | <0.001 |
| CBCL Total T Score | 61.57(11.86) | 65.51(8.81) | 65.28(9.71) | 62.21(8.71) | 44.23(9.37) | 33.5 | <0.001 |
| CBCL Externalizing Behavior T Score | 56.33(12.17) | 59.32(10.74) | 60.68(11.44) | 54.23(10.81) | 42.94(7.86) | 17.52 | <0.001 |
| CBCL Internalizing Behavior T Score | 61.45(11.4) | 64.5(9.7) | 61.98(10.83) | 65.54(9.47) | 48.59(9.33) | 17.9 | <0.001 |
| Full Scale IQ (age-dependent) | 100.9(19.24) | 95.31(21.32) | 100.9(14.83) | 112.7(21.88) | 110.9(10.46) | 9.41 | <0.001 |
| rs-fMRI SAMPLE |  |  |  |  |  |  |  |
|  | Total | ASD | ADHD | OCD | TDC | X<sup>2</sup> | p-value |
| N | 299 | 113 | 85 | 50 | 51 |  |  |
| Males | 214 | 88 | 61 | 31 | 34 | 4.04 | 0.26 |
| DWI SAMPLE |  |  |  |  |  |  |  |
| N | 157 | 78 | 38 | 31 | 10 |  |  |
| Males | 119 | 62 | 32 | 22 | 3 | 13.87 | 0.003 |

Note: ASD = Autism Spectrum Disorder, ADHD = Attention-Deficit/Hyperactivity Disorder, OCD = Obsessive Compulsive Disorder, TDC = Typically Developing Children. This table shares the CBCL T-scores which are normalized for age and gender. The analyses used CBCL raw scores (not normalized for age and gender).

### Brain/Behavior-age relations

Linear regression analyses between age, age-squared, and brain and behavior indices confirmed that relations between these indices of interest and age were mainly linear and consistent with the prior literature (see supplementary; *Table S2, Figure S6*).

### Cortico-amygdalar structural covariance

No significant interaction effects between either internalizing or externalizing behavior score and left or right amygdala volume on vertex-wise cortical thickness were found. No significant effects were found when diagnostic status was included into the interaction term (i.e., diagnostic status-by-externalizing/internalizing behavior-by-left/right amygdala volume on cortical thickness). *Figure 2* illustrates the unthresholded p-maps of the relationship between externalizing and internalizing behavior and left cortico-amygdalar structural covariance/functional connectivity.

**Figure 2.**
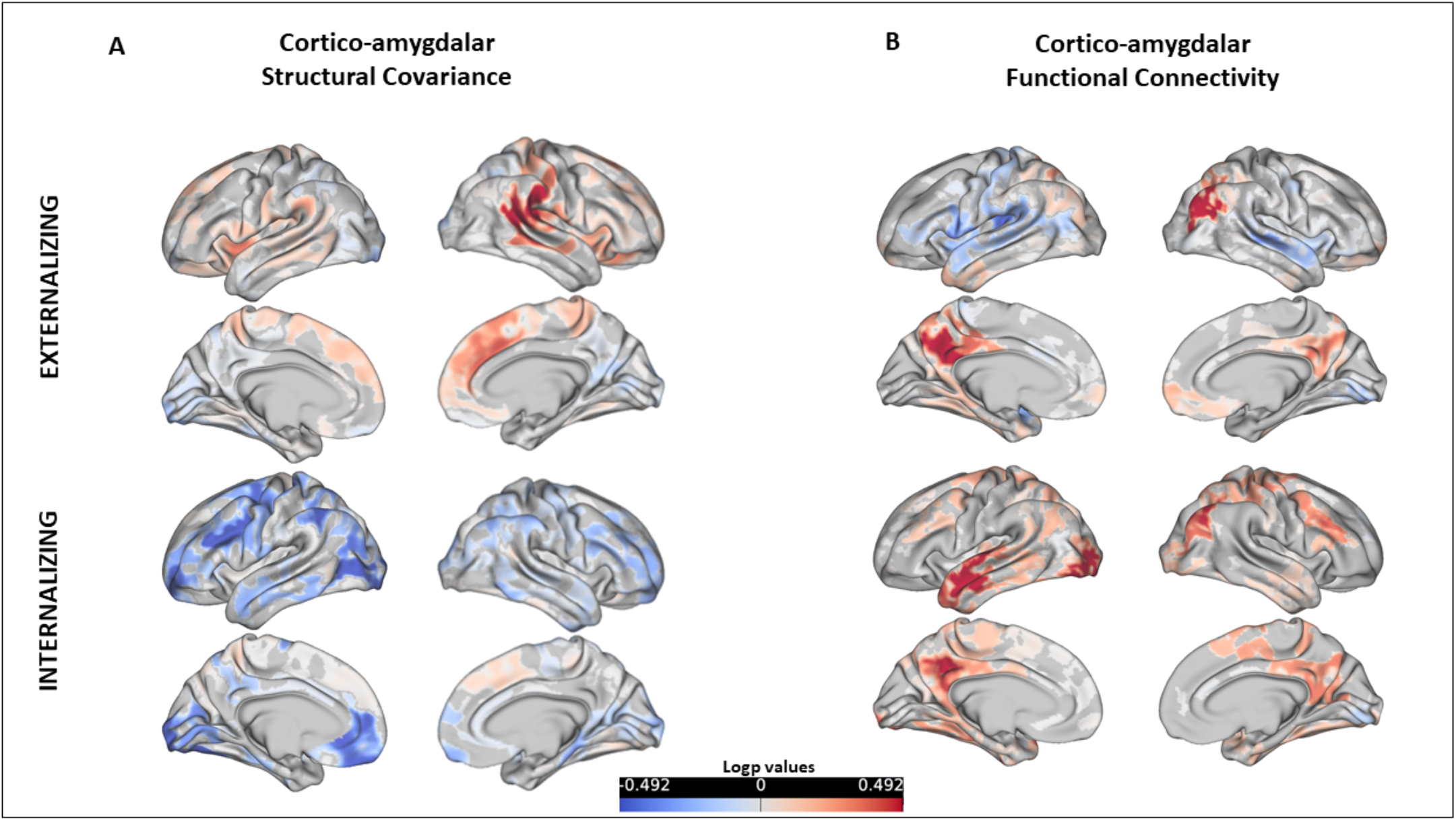
Unthresholded spatial p-map of the relationship between externalizing/internalizing behaviors and cortico-amygdalar structural and functional connectivity. A) Unthresholded spatial p-maps depicting the relationship between the interaction of externalizing and internalizing behavior and left amygdala volume on each cortical vertex. B) Unthresholded spatial p-maps depicting the relationship between externalizing and internalizing behavior and functional connectivity between the left amygdala seed and each cortical vertex. A logp value of 1.6 is considered significant. As seen in the figure, none of the results reached this significance threshold.

### Functional and white matter connectivity

No significant associations were found between either externalizing or internalizing behavior and the time series correlations between left or right amygdala volume and each cortical vertex (*Figure 2*) and no significant interaction effect between externalizing or internalizing and diagnostic status on functional connectivity. Similarly, there were no significant associations found between either behavioral domain and FA or MD of the left or right UF or CB (*Table 2, Figure 3*). No diagnosis-by-behavior interaction effects were found across the functional and white matter connectivity models.

**Table 2.** Linear Model Results of the white matter connectivity analysis

| White Matter Tract | Behavioral Variable | Beta | 95% CI (X, X) | t<sub>(2,153)</sub> | p-value |
| --- | --- | --- | --- | --- | --- |
| Left UF FA | Externalizing Behavior | -0.005 | (-0.0006, 0.00058) | -0.062 | 0.95 |
|  | Internalizing Behavior | 0.092 | (-0.00028, 0.00103) | 1.13 | 0.261 |
| Left UF MD | Externalizing Behavior | 0.043 | (-4.61e-07, 8.12e-07) | 0.546 | 0.59 |
|  | Internalizing Behavior | -0.046 | (-9.01e-07, 4.92e-07) | -0.58 | 0.562 |
| Right UF FA | Externalizing Behavior | 0.012 | (-0.00047, 0.00054) | 0.131 | 0.896 |
|  | Internalizing Behavior | 0.009 | (-0.0005, 0.00058) | 0.105 | 0.917 |
| Right UF MD | Externalizing Behavior | 0.139 | (-4.55e-08, 1.09e-06) | 1.818 | 0.071 |
|  | Internalizing Behavior | 0.024 | (-5.34e-07, 7.29e-07) | 0.304 | 0.761 |
| Left CB FA | Externalizing Behavior | -0.009 | (-0.00056, 0.00049) | -0.13 | 0.897 |
|  | Internalizing Behavior | -0.039 | (-0.00073, 0.00042) | -0.53 | 0.597 |
| Left CB MD | Externalizing Behavior | 0.109 | (-7.5e-08, 6.95e-07) | 1.592 | 0.113 |
|  | Internalizing Behavior | 0.028 | (-3.3e-07, 5.13e-07) | 0.405 | 0.68 |
| Right CB FA | Externalizing Behavior | -0.003 | (-0.0005, 0.0005) | -0.041 | 0.967 |
|  | Internalizing Behavior | -0.039 | (-0.00074, 0.00042) | -0.53 | 0.597 |
| Right CB MD | Externalizing Behavior | 0.109 | (-7.49e-08, 6.97e-07) | 1.592 | 0.113 |
|  | Internalizing Behavior | 0.028 | (-3.4e-07, 5.1e-07) | 0.405 | 0.69 |

Note: UF = Uncinate fasciculus, CB = Cingulum Bundle, FA=Fractional Anisotropy, MD = Mean Diffusivity.

**Figure 3.**
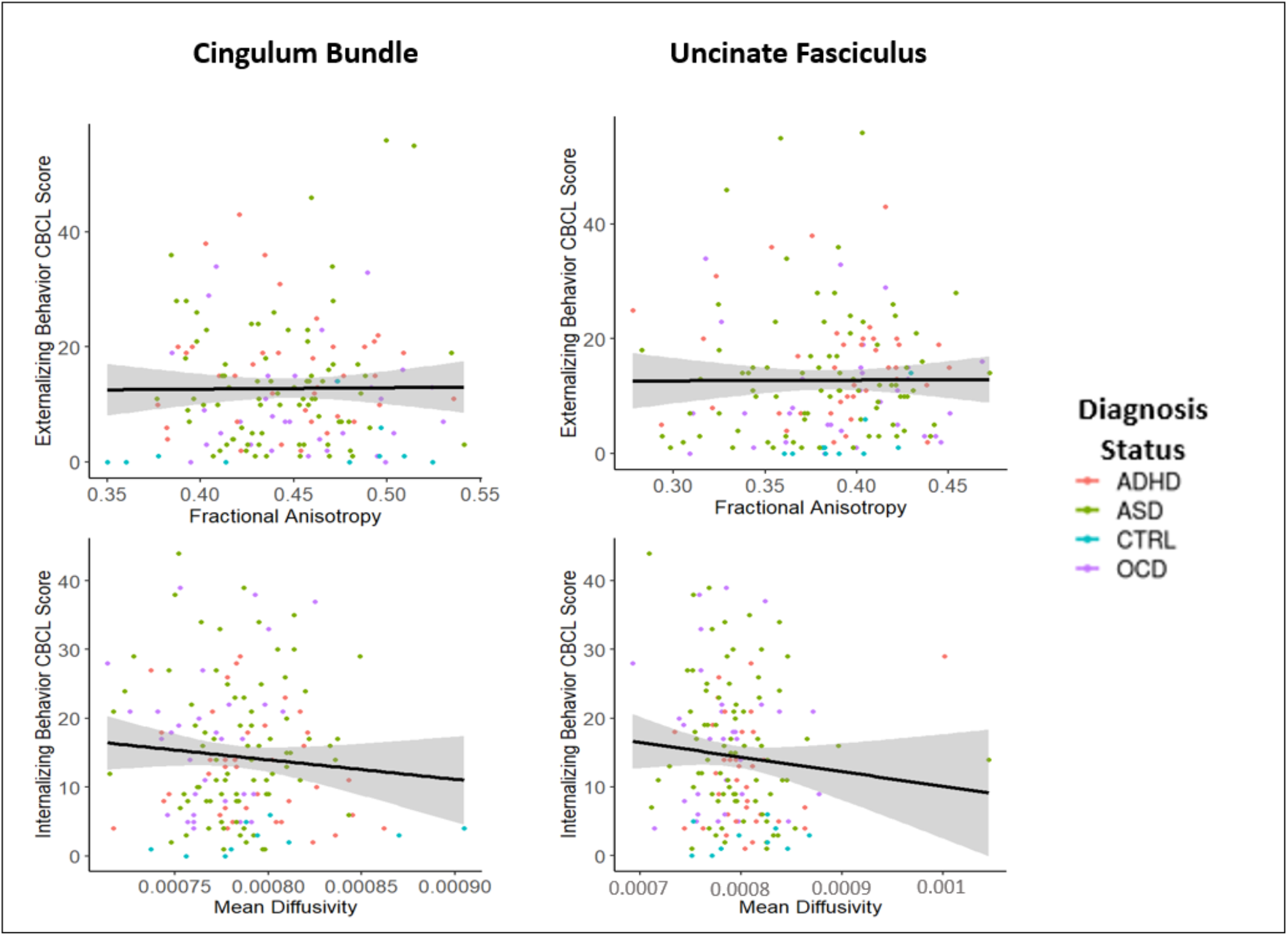
Relationship between externalizing or internalizing behavior and fractional anisotropy and mean diffusivity (units: mm^2^/s) of the two white matter tracts of interest: the cingulum bundle and uncinate fasciculus. The depicted relationships are all non-significant. The black line is the regression line and the shaded gray area is the confidence interval. These figures include all data points, including potential outliers. Analyses were run with and without outlier removal; the results remained non-significant in either case. ADHD = Attention Deficit Hyperactivity Disorder, ASD = Autism Spectrum Disorder, OCD = Obsessive Compulsive Disorder, CTRL = healthy control/typically developing

### Planned Subsample Analysis

Findings remained the same for the structural covariance and functional connectivity analyses among the subset of participants with a time-gap of one month or less between imaging acquisition and behavioral assessments. Among the white matter connectivity models, there was a significant association between MD of the right UF and externalizing behavior, however, this association no longer remained when participants with outlier MD values were removed (*Figure S7*). There was a main effect of externalizing behavior on the MD of the right CB, such that higher MD was associated with greater externalizing behavior (*Figure S8*) which did not survive the FDR significance threshold (F_3,106_=16.6, p_Model_ <0.001, t_Externalizing_= 2.19, p_Externalizing_=0.03). There were no other significant associations found between FA or MD of the left or right UF or CB.

### Post-hoc bootstrap resampling analysis

See *Figure 4* for structural covariance and functional connectivity bootstrap resampling analyses (and *Figure S9* for other models) plotting the regression coefficients and their standard errors at each vertex (Panel A). Across all models, the regression coefficients based on 1000 generated bootstrapped resampled models were near zero (<0.01; low signal) with respective standard errors near zero (<0.001; low noise). Vertices with a t-statistic greater than an absolute value of 4 served as a proxy for a high signal vertex and are colored in pink in *Figure 4*. This proxy serves to provide insight into the stability of both high and low signal vertices. The figure illustrates both high and low signal vertices have low standard errors of the regression coefficients (<0.001) indicating a stable zero. Panels B-D of *Figure 4* illustrates the distribution of three mean model parameter estimates (averaged across the 1000 iterations); regression coefficients (structural covariance: 2.56e-06 ± 1.37e-05; functional connectivity: 0.0002 ± 0.004), t-statistics (structural covariance: 0.357 ± 1.91; functional connectivity: 0.119 ± 1.99), and effect sizes (structural covariance: 8.45e-06 ± 4.66e-05; functional connectivity: 0.00045 ± 0.009). For the white matter connectivity models, the bootstrapped standard errors of the regression coefficients were also near zero and nearly all models featured bootstrapped confidence intervals which included zero (*Table S5*). See *Figure S10* and *Table S6* for bootstrap resampled results for the sensitivity analyses.

**Figure 4.**
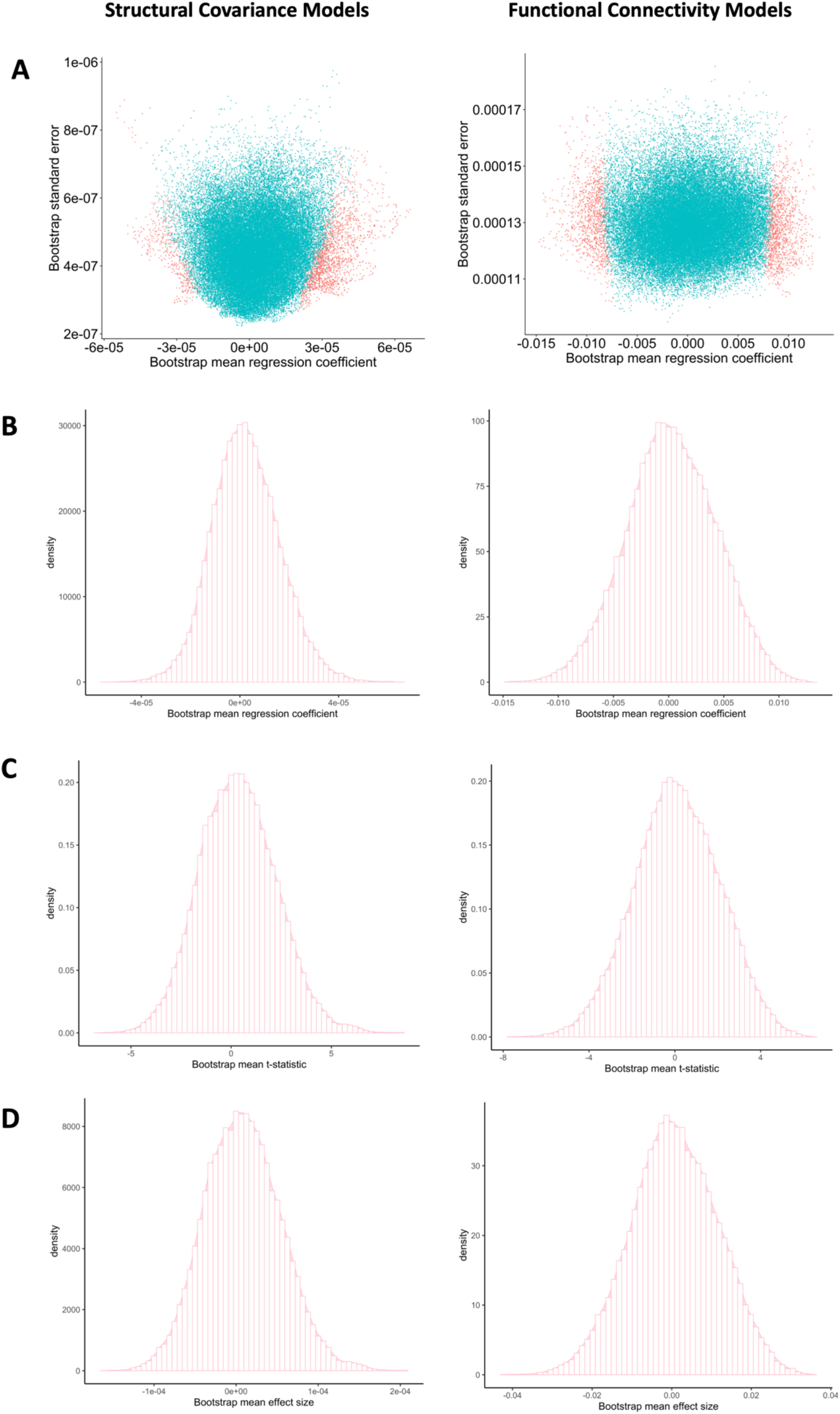
Bootstrap resampling of the externalizing behavior*left amygdala structural covariance and functional connectivity models. All other models examined feature similar relationships between standard errors and regression coefficients (see supplementary). This figure depicts the results of a bootstrap resampling analysis in which 1000 iterations of each data matrix were generated and used to perform the linear regression models in PALM. Panel A illustrates scatterplots between the mean regression coefficient (averaged across 1000 resamples) and the bootstrapped standard errors of the regression coefficients of each vertex for the structural covariance and functional connectivity models. Points colored in pink depict the vertices with a t-statistic greater than 4 to illustrate a proxy for ‘higher signal’ and low noise vertices. Points colored in blue depict the vertices with a t-statistic less than 4 to illustrate ‘low signal’ and low noise vertices. Panel B depicts the histogram of the mean regression coefficients of each vertex across the 1000 bootstrapped resampled analyses. Panel C depicts the histogram of the mean t-statistic of each vertex across the 1000 bootstrapped resampled analyses. Panel D depicts the histogram of the mean effect size of the model at each vertex across the 1000 bootstrapped resampled analyses.

## DISCUSSION

Using a multi-modal imaging framework, the current study did not find a continuous (dimensional) linear relationship between cortico-amygalar connectivity properties and externalizing or internalizing behaviors across a transdiagnostic sample including TDC and children with ASD, ADHD, or OCD. Further, our results did not suggest that the brain-behavior patterns examined differed by diagnostic group. Importantly, post-hoc bootstrap resampling results indicate reliability and stability of the null results found in the present study.

The previous neuroimaging studies that found a significant linear association between internalizing or externalizing behavior and cortico-amygdalar connectivity metrics included mainly TDC samples and mostly analyzed structural MRI metrics (Albaugh et al., 2016; Ameis et al., 2014; Andre et al., 2019; Ducharme et al., 2011; Mohamed Ali et al., 2019; Vijayakumar et al., 2017). Among these, studies utilizing moderate-to-large (as opposed to smaller) sample sizes reported small effect sizes for associations found (*see Table S3*). The current study featured a moderate-to-large sample size, indicating that we likely had sufficient power to detect a small effect size of the linear brain-behavior relationship of interest. The near-zero standard errors of regression coefficients for the main models submitted to bootstrap resampling (found across low and high signal vertices), coupled with relatively low standard deviations of model parameter estimates, suggests that the null results found in the current study are stable and reliable. Our transdiagnostic NDD sample likely exhibited greater behavioral (and potentially brain-level) heterogeneity than prior positive studies in TDC (Dajani et al., 2016; Dickie et al., 2018; Fair et al., 2012). The stability of our null results may indicate that univariate statistical approaches may not be optimal to find shared brain-behavior relations within large transdiagnostic NDD samples (as seen in other reports, e.g., Marek et al. 2020). The small effect sizes found across the bootstrapped resampled structural covariance and functional connectivity models also suggests that the effect size of any linear relationship between externalizing or internalizing behavior and cortico-amygdalar connectivity properties is either not present or too small to be detected across a heterogeneous transdiagnostic clinical sample of children with different NDDs.

Although the current study examined cortico-amygdalar connectivity across three imaging modalities, it is important to note that it only assessed behavioral traits through a single broad-band parent-report measure. Delineating brain-behavior relations relevant to internalizing or externalizing behaviors in heterogeneous clinical samples may benefit from incorporating multi-modal measures of behavior (e.g., task-based fMRI using behavioral relevant tasks). Multi-modal measures of behavior may improve the estimate of the trait(s) of interest compared to use of a parent-report behavioral measure alone. Two previous studies have examined brain-behavior relationships in transdiagnostic samples using both symptom measures and task-based fMRI; Ibrahim et al. found a negative association between cortico-amygdalar connectivity during an emotion perception task and externalizing behavior score across children with ASD, with or without co-occurring disruptive behavior disorders (Ibrahim et al., 2019). Stoddard et al. found that amygdala-prefrontal cortex connectivity during viewing of intensely angry faces was associated with different behavioral profiles across a sample of children with ADHD, disruptive behavior disorders, anxiety disorders or TDC, whereby decreased connectivity was associated with high levels of anxiety and irritability, and lower connectivity was associated with high anxiety but low irritability (Stoddard et al., 2017).

Given the potential for differing brain-behavior profiles across a heterogeneous clinical group (i.e., one or more NDD populations may include subgroups with differing underlying distributions (Fair et al., 2012)), the use of clustering approaches may be an effective technique to parse through this heterogeneity and distinguish the boundaries of different brain-behavior profiles between NDD subgroups (Feczko et al., 2019). Clustering has recently been applied to the POND dataset and has identified transdiagnostic subgroups of children with different NDD diagnoses that exhibit more similar (within-subgroup) brain-behavior profiles and larger between-subgroup differences than those found based on DSM category (Jacobs et al., 2020; Kushki et al., 2019). One of these studies included CBCL-externalizing and internalizing behaviors as features in the clustering model and found different distributions exhibited among newly identified subgroups for externalizing and internalizing behaviors, though neither of these features contributed significantly to the cluster solution (Jacobs et al., 2020).

A number of strengths and limitations of the current study require consideration. First, considering the growing concern of statistical practices which may contribute to false positive results or inflated effect sizes (Marek et al., 2020; Poldrack et al., 2017), we made use of non-parametric statistics (Eklund et al., 2016) (i.e. TFCE; (Smith and Nichols, 1996)) to reduce risk of inflation for any potential associations found. Additionally, a standardized QC protocol was implemented across all imaging modalities to reduce the likelihood for findings to be driven by artefacts or motion (Backhausen et al., 2016; Pardoe et al., 2016). This resulted in an exclusion rate of 16.8-23.7% across imaging modalities based on image QC (that is, following initial exclusion of participants based on age, missing data and >12-month time-gap between imaging and behavioral assessments), which is comparable to previous studies examining pediatric samples or using standardized QC approaches (Ameis et al., 2014; Ducharme et al., 2014; Xia et al., 2018) and in line with higher in-scanner motion in pediatric and clinical samples (Pardoe et al., 2016). While applying this rigorous QC approach is beneficial (particularly in a pediatric clinical sample), this limited our ability to leverage more of the total data available from POND. The rs-fMRI (n=299) and DWI (n=157) samples were smaller than the T1-weighted sample (n=346), further reducing statistical power. Further, T1-weighted and rs-fMRI acquisitions for this sample were collected across a scanner upgrade, potentially introducing scanner-related confounds not captured by our statistical approaches. Lastly, while parent-report behavioral measures of internalizing or externalizing behaviors have been used in prior brain-behavior studies (Albaugh et al., 2016; Ameis et al., 2014; Ducharme et al., 2014, 2011; Ibrahim et al., 2019), inclusion of additional measures (e.g., self-report behavioral or cognitive) may provide a more sensitive proxy of the behavioral domain of interest than use of a parent-report measure alone to relate with brain indices.

## CONCLUSION

Producing consistent results that are generalizable and replicable has been challenging in clinical and cognitive neuroscience (Ioannidis, 2018; Simmons et al., 2011) as suggested by reports of non-replication (He et al., 2020), heterogeneous findings (Uddin et al., 2017), and negative findings (Dajani et al., 2019; Masouleh et al., 2019). Null reports are necessary to refine methodological approaches which can inform future research. Contrary to our hypotheses, the stability and reliability of the null result found in the current study (across three imaging modalities) provides support for the absence of a dimensional linear association between externalizing or internalizing behavior and cortico-amygdalar connectivity across a heterogeneous group of children with different NDD diagnoses and TDCs. Future work exploring brain-behavior relations relevant to internalizing and externalizing domains in transdiagnostic samples may benefit from the use of additional clinical/cognitive/behavioral assessments (including multi-informant reports or relevant task-based fMRI), and clustering approaches to delineate subgroups with different brain-behavior profiles.

## Acknowledgements

We thank the following individuals for research support and data collection: Tara Goodale, M.Sc., Reva Schachter, M.Sc., Mithula Sriskandarajah, B.Sc., Marlena Colasanto, M.Sc., Jennifer Gomez, M.A., and Laura Park, M.Sc, from The Hospital for Sick Children; Susan Day Fragiadakis, M.A., Naomi Peleg, M.Sc., and Leanne Ristic, B.A., from Holland Bloorview Kids Rehabilitation Hospital; Richa Mehta, B.A., Christina Sommerdyk, M.Sc., from the Lawson Health Research Institute; Carolyn Russell, B.Sc., Alessia Greco, M.A., Mike Chalupka, B.A., B.Sc., Christina Chrysler, B.A., Irene O’Connor, M.Ed. Psych., from McMaster Children’s Hospital, Melissa Hudson from Queen’s University. A special thank you to Alana Iaboni and Christopher Hammill for their extraordinary support in clarifying questions related to behavioral and imaging data, respectively, in the POND sample. An additional special thank you to Dr. Anthony Randal McIntosh who provided assistance with constructing and interpreting the bootstrap resampling analysis.

## Funding, Disclosures and conflict of interest

This research was supported by the grant IDS-I l-02 from the Ontario Brain Institute. The Ontario Brain Institute is an independent non-profit corporation funded partially by the Ontario government. The opinions, results and conclusions are those of the authors and no endorsement by the Ontario Brain Institute is intended or should be inferred. HN has received funding from the CAMH Discovery Fund and currently receives funding from the Ontario Graduate Scholarship. GRJ received funding from the Ontario Graduate Scholarship and Ontario Student Opportunity Trust Fund. ANV currently receives funding from the National Institute of Mental Health (1/3R01MH102324 & 1/5R01MH114970), Canadian Institutes of Health Research, Canada Foundation for Innovation, CAMH Foundation, and University of Toronto. NJF received funding from the Centre for Addiction and Mental Health Discovery Fund Postdoctoral Grant. M-CL receives funding from the Ontario Brain Institute via the POND Network, Canadian Institutes of Health Research, the Academic Scholars Award from the Department of Psychiatry, University of Toronto, and CAMH Foundation. PS has received royalties from Guilford Press. RS has consulted to Highland Therapeutics, Eli Lilly and Co., and Purdue Pharma. He has commercial interest in a cognitive rehabilitation software company, “eHave”. PDA receives funding from the Alberta Innovates Translational Health Chair in Child and Youth Mental Health and holds a patent for ‘SLCIAI Marker for Anxiety Disorder’ granted May 6, 2008. EA receives funding from the Canadian Institutes of Health Research, National Institutes of Health, Ontario Brain Institute, Brain Canada, Azrieli foundation, Autism Speaks, Health Resources & Services Administration. She has served as a consultant to Roche and Quadrant, has received grant funding from Roche, holds a patent for the device, “Anxiety Meter”, has received editorial honoria from Wiley and royalties from APPI and Springer. SHA currently receives funding from the National Institute of Mental Health (R01MH114879), Canadian Institutes of Health Research, the Academic Scholars Award from the Department of Psychiatry, University of Toronto, Autism Speaks and the CAMH Foundation. Other authors report no related funding support, financial or potential conflicts of interest.

## Notes

https://github.com/hajernakua/cortico-amygdalar2019

