## Supplementary Materials for "Cortico-amygdalar connectivity and externalizing/internalizing behavior in children with neurodevelopmental disorders"

**Table of Contents**

Quality Control ……………………………………………………………………………..2

Preprocessing Pipelines……………………………………………………………………..4

Further Processing and Analysis Details……………………………………………………5

Sample Characteristics………………………………………………………………………6

Baseline Functional and White Matter Connectivity Regression Models.………………………..8

Comparing relationship between brain and behavioral metrics and age with previous reports….9

Post-hoc Bootstrap Resampling…………………………………………………….……..10

Tables……………………………………………………………………………………11

Figures…………………………………………………………………………………..17

References……………………………………………………………………………….30

### 1. Quality Control (QC)

Recent research has suggested that quality control (QC) decisions influence neuroimaging analyses and outcomes (1–3). As a result, careful considerations were employed when deciding the QC procedure for the current study.

#### *1.1. T1-weighted Structural Images*

This QC procedure incorporates qualitative (visual) and quantitative (image quality metrics) assessments of quality. Visual QC was carried out through visual inspection of the QC HTML pages generated by the fMRIprep pipeline (version 1.1.1; completed in 2018). Each scan was rated on a scale of 1-5 based on the severity of the visible artefacts and its influence on cortical segmentation, with 1 being the best quality and 5 being the worst (*Figure S1*). Quantitative QC involved use of image quality metrics (IQMs) produced by the MRIQC pipeline (4) with three metrics of interest: contrast-to-noise ratio (CNR), signal-to-noise ratio (SNR) and coefficient of joint variation (CJV). These three metrics were chosen as they provide information on scan quality and are related to in-scanner head motion. Quantitative metrics may catch subtleties missed by visual QC.

During visual QC, each scan was evaluated based on: (1) whether the pipeline correctly extracted the brain from the skull (if there was incorrect extraction, the pipeline was re-run, and if there was persistent incorrect extraction, that participant would be excluded), (2) the quality of the T1 image and template (e.g. artefacts or truncated brain regions could be detected), and (3) segmentation of the pial and white matter surface layer. If a participant had segmentation inaccuracies as a result of processing issues, their brain scan was re-processed and underwent the QC procedure once again. However, if the segmentation was incorrect due to a poor quality T1 acquisition (i.e. noticeable artefacts), then those participants were excluded. After visual inspection and rating, participants were placed in one of three groups: pass (score of 1 or 2), uncertain (score of 3) or fail (score of 4 or 5). The QC threshold implemented in this study excluded participants with a score of 4 or 5. In addition to the visual QC procedure, subjects who had a CNR, SNR and/or CJV value greater than 2 standard deviations from the mean were also excluded. The protocol for the QC procedure can be found here (https://github.com/hajernakua/cortico-amygdalar2019).

Automated QC decisions from the MRIQC random forest classifier (4) were used in place of a second rater. This classifier, which was trained on the ABIDE and DS030 datasets, predicts whether a subject should be accepted or rejected based on 14 image quality metrics. Using the 'irr' package in R (version 3.5.0), an interrater reliability analysis was run to compare rates of inclusion and exclusion between the standard QC level and the random forest classifier. The results revealed a kappa value of 0.591 (*kappa2* function) with a 90.5% agreement (*agree* function) between the random forest classifier and the pass and fail criteria.

*1.2. Resting-state fMRI*

Following data processing and cleaning, mean framewise displacement (FD) from the MRIQC pipeline was used as a criterion for exclusion of resting-state fMRI data (5–7). Due to the nature of the sample being children with psychiatric conditions who are reported to have greater in-scanner head motion (7–9), we used a mean FD >0.5mm as our threshold of exclusion.

*1.3. Diffusion Weighted Images*

Processed diffusion data were examined for artefacts and processing errors via qualitative and quantitative QC. Visual inspection assessed slices from each orthogonal view of the B0, brain mask, first eigenvector (V1/FA direction) and residuals from a tensor model fit via DTIFIT. Participants were excluded based on indications of slice dropout, inter-slice variability, and truncated brain regions. Quantitative QC excluded participants with CNR, SNR, relative motion and number of outlier slice values greater than two standard deviations from the mean. Participants who failed either qualitative or quantitative QC were excluded from analyses. Further details are included in the DWI QC protocol (https://github.com/hajernakua/cortico-amygdalar2019).

### 2. Pre-processing pipelines

#### *2.1. Structural Images*

Copied from the fMRIPrep readthedocs: Results included in this manuscript come from preprocessing performed using FMRIPREP version 1.1.1 [1, 2, RRID:SCR_016216], a Nipype [3, 4, RRID:SCR_002502] based tool. Each T1w (T1-weighted) volume was corrected for INU (intensity non-uniformity) using N4BiasFieldCorrection v2.1.0 [5] and skull-stripped using antsBrainExtraction.sh v2.1.0 (using the OASIS template). Spatial normalization to the ICBM 152 Nonlinear Asymmetrical template version 2009c [7, RRID:SCR_008796] was performed through nonlinear registration with the antsRegistration tool of ANTs v2.1.0 [8, RRID:SCR_004757], using brain-extracted versions of both T1w volume and template. Brain tissue segmentation of cerebrospinal fluid (CSF), white-matter (WM) and gray-matter (GM) was performed on the brain-extracted T1w using fast [17] (FSL v5.0.9, RRID:SCR_002823).

#### *2.2. Resting-state Functional Images*

Copied from the fMRIprep readthedocs: Functional data was slice time corrected using 3dTshift from AFNI v16.2.07 [11, RRID:SCR_005927] and motion corrected using mcflirt (FSL v5.0.9 [9]). This was followed by co-registration to the corresponding T1w using boundary-based registration [16] with 9 degrees of freedom, using flirt (FSL). Motion correcting transformations, BOLD-to-T1w transformation and T1w-to-template (MNI) warp were concatenated and applied in a single step using antsApplyTransforms (ANTs v2.1.0) using Lanczos interpolation.

Physiological noise regressors were extracted applying CompCor [18]. Principal components were estimated for the two CompCor variants: temporal (tCompCor) and anatomical (aCompCor). A mask to exclude signal with cortical origin was obtained by eroding the brain mask, ensuring it only contained subcortical structures. Six tCompCor components were then calculated including only the top 5% variable voxels within that subcortical mask. For aCompCor, six components were calculated within the intersection of the subcortical mask and the union of CSF and WM masks calculated in T1w space, after their projection to the native space of each functional run. Framewise displacement [19] was calculated for each functional run using the implementation of Nipype.

Many internal operations of FMRIPREP use Nilearn [22, RRID:SCR_001362], principally within the BOLD-processing workflow. For more details of the pipeline see https://fmriprep.readthedocs.io/en/latest/workflows.html.

### 3. Further Processing and Analysis Details

#### *3.1. Using PALM for structural and functional analyses*

The general linear models used for the vertex-wise cortical structural covariance and functional connectivity analyses were implemented using FSL's permutation analysis of linear models (PALM) tool. We examined cortico-amygdalar structural covariance and seed-based functional connectivity at the group level. The analysis used 2000 permutations with threshold free cluster enhancement (TFCE) as implemented in FSL's PALM. Group results were thresholded at $p<0.05$ FDR-corrected for the number of vertices in each hemisphere, and further corrected for separate runs of PALM for each hemisphere (i.e. the critical TFCE-corrected threshold was set to $p<0.025$) (10).

#### *3.2. Deterministic Tractography with Slicer dMRI*

Once pre-processing and QC of the diffusion data were completed, the Slicer dMRI software was used to perform deterministic tractography. A study-specific atlas was created which used the tractography fibers from 21 subjects who captured the variability of the dataset (based on age, gender, and diagnosis) and had good quality dMRI acquisitions. Slicer organizes individual fibre streamlines into clusters which were then used to manually form the white matter tracts based on visual guidance from the Catani white matter atlas (11). From there, the atlas was registered to each participant, and diffusion metrics (i.e. fractional anisotropy and mean diffusivity) were extracted and analyzed. Prior to analysis, quality of streamlines (visually and quantitatively) were assessed. Output HTML files from Slicer were visually assessed based on missing streamlines or errors in registration of the tracts at the participant level. Streamlines and tracts were quantitatively assessed by determining the range and mean of streamline fibers for all analyzed tracts across participants.

*3.2.1 Overall noise as a covariate for the diffusion analysis*

Once preprocessing was completed, residual noise was estimated using MRtrix's *dwidenoise* function. Average noise was estimated across DWI scans at every voxel of each participant's brain acquisition. These were summarized into a mean score per participant (after removing missing values). This variable was used as a covariate if the baseline regression model examining white matter connectivity metrics were significant.

### 4. Sample Characteristics

*4.1. T1-weighted Imaging Sample*

There were no significant differences in externalizing (t=0.06, p=0.95) and internalizing (t=0.74, p=0.46) behavior scores between the total POND sample (n=611) and the present study sample (n=346), nor were there any differences in age (t=0.63, p=0.53), gender ($X^2$=0.58, p=0.75), or diagnostic composition ($X^2$=4.74, p=0.31). There was a significant correlation between age and internalizing behavior (r=0.17, p<0.001). There was no significant difference between included and excluded participants across the scanner upgrade ($X^2$=0.13, p=0.72). There were 16 participants in the T1w sample who had an IQ < 70 (mean= 52.5, range=40-67).

*4.1.1. Medication*

Of the 346 participants included in the T1w analysis, 146 (42.2%) of them were currently on or had been on a prescription medication in the past 6 months. A total of 160 (46.2%) reported no medication and 40 (11.6%) did not have medication information available. Among the diagnostic groups, 65 (46.4%) with ASD, 40 (40%) with ADHD and 17 (32.1%) with OCD reported being on medications. In the subset of the sample with functional (n=299) and diffusion data (n=157), 126 (43.6%) and 67 (42.7%) reported taking medication within the past 6 months, respectively.

*4.2. rs-fMRI Sample*

There were no significant differences in externalizing (t=0.04, p=0.96) and internalizing (t=0.6, p=0.55) behavior scores between the total rs-fMRI POND sample (n=574) and the present study sample (n=299), nor were there any differences in age (t=0.58, p=0.56), gender ($X^2$=0.86, p=0.65), or diagnostic composition ($X^2$=6.39, p=0.17). Within the main rs-fMRI sample (n=299), age was significantly correlated with internalizing (r=0.15, p=0.009) but not externalizing (r=-0.11, p=0.06) behaviors. Motion, as indexed by mean FD, was significantly correlated with externalizing (r=0.12, p=0.039) but not with internalizing behavior (r=-0.05, p=0.35). There was no significant difference in mean FD between the scanners (t=-0.56, p=0.58). There were 15 participants in the rs-fMRI sample who had an IQ < 70 (mean= 52.7, range=40-68).

*4.3. Diffusion Weighted Imaging Sample*

There were no significant differences between age (t=-0.54, p=0.59) externalizing (t=0.47, p=0.63) and internalizing (t=-0.36, p=0.72) behavior between the fully acquired DWI sample (n=262) and the analyzed sample (n=157). In the analyzed sample, age was correlated with internalizing behavior (r=0.22, p=0.005), but not externalizing behavior (r=-0.02, p=0.79). Overall noise was not correlated with either externalizing (r=0.02, p=0.8) or internalizing (r=0.12, p=0.13) behavior. There were 7 participants in the DWI sample who had an IQ < 70 (mean= 52.4, range=40-66).

*4.4. Subsample – T1-weighted Imaging Acquisitions*

The subsample analysis excluded 94 additional participants from the main sample resulting in 252 participants analyzed (*Figure S2*). There were no significant differences in age (t=-0.2, p=0.84), externalizing (t=0.67, p =0.49) and internalizing (t=0.06, p=0.95) behavior between the main cohort analysed (n=346) and this sub-sample (n=252). There was a significant difference in diagnosis composition across the scanner upgrade ($X^2$=39.4, p<0.001). There were 14 participants in the T1w subsample who had an IQ < 70 (mean= 50.8, range=40-67).

*4.5. Subsample – Resting state fMRI Acquisitions*

For the rs-fMRI subsample analysis, we analyzed 217 participants. There were no significant differences between externalizing (t=1.01, p=0.31) and internalizing (t=0.67, p=0.49) behavior, age (t=0.16, p=0.87), or mean FD (t=-0.47,p=0.64) between the main sample (n=299) and sub-sample (n=217), nor were there differences in gender ($X^2$=0.04, p=0.83) and diagnosis

($X^2$=2.05, 0.56) composition. There were 13 participants in the sensitivity rs-fMRI sample who had an IQ < 70 (mean= 52.9, range=40-68).

*4.6. Subsample – Diffusion Weighted Imaging*

There were no significant differences between age (t=0.51, p=0.61) externalizing (t=-0.17, p=0.86) and internalizing (t=0.54, p=0.58) behavior between the main analyzed sample (n=157) and the subsample cohort (n=110). Further, there were no differences in the gender ($X^2$=0.18, p=0.67) or clinical diagnosis ($X^2$=6.53, p=0.09) composition between these two cohorts. In the subsample (n=110), age was correlated with internalizing behavior (r=0.24, p=0.012), but not externalizing behavior (r=-0.02, p=0.82). Overall noise was correlated with internalizing (r=0.19, p=0.048) but not externalizing (r=0.06, p=0.51) behavior. There were 5 participants in the DWI subsample who had an IQ < 70 (mean= 52.2, range=42-60).

**5. Baseline Functional and White Matter Connectivity Regression Models**

*Functional Connectivity Left or Right = Intercept +* $\beta_1$*(externalizing or internalizing behavior CBCL score) +* $\beta_2$*(age) +* $\beta_3$*(gender) +* $\beta_4$*(scanner) +* $e_j$*.*

*FA/MD (Left or Right Uncinate Fasciculus/Cingulum Bundle) = Intercept +* $\beta_1$*(externalizing or internalizing behavior CBCL score) +* $\beta_2$*(age) +* $\beta_3$*(gender) +* $e_j$*.*

*5.1. Including the Age Term*

Prior to fitting the structural covariance, functional connectivity, and white matter connectivity models, the relationship between the brain regions of interest and age and age-squared were examined. The better fitting age term was included as the covariate in the main analyses (*Table S2*). If the better fitting age term was quadratic, then the linear and quadratic age terms were included in the model. RStudio and PALM were used to model the age-brain metric relationship for structural covariance/white matter connectivity and functional connectivity, respectively. Age or age-squared was included as the dependent variable with left or right cortical thickness, amygdala volume, diffusion metrics, or seed-based functional connectivity between amygdala and cortical vertices as the independent variable. Age nor age-squared was significantly associated with FA of the left UF, although there was a slightly better fit with age-squared. Models with FA of the left UF as the dependent variable were separately fit with age, age + age-squared and no age variable to ensure that these variables did not influence the brain-behavior relationship of interest. Including any of these age terms did not influence the relationship between externalizing or internalizing behavior and FA of the left UF. All other structural covariance and white matter connectivity analyses included age as a linear term. Similarly, using PALM, age nor age-squared were significantly associated with seed-based functional connectivity. All functional connectivity models were run with age included as a linear term and again without age included in the model. The results remained the same.

**6. Comparing relationship between structural brain and behavioral metrics and age with previous reports**

To rule out that the null results of this study were due to a noisy or atypical NDD dataset, we examined the relationships between age and: (i) cortical thickness (indexed using mean left and right cortical thickness based on Desikan-Killiany Atlas parcellations), (ii) left and right amygdala volume (based on Desikan-Killiany Atlas parcellations), (iii) FA and MD of the uncinate fasciculus (UF) and cingulum bundle (CB), and (iv) externalizing and internalizing behaviors. We examined this relationship across the analyzed samples (n=346 or n=157) (*Table S2; Figure S6a-c*).

Within the sample, both left and right cortical thickness (12) and amygdala volume followed a linear trajectory across age (13). Although some reports suggest a quadratic relationship between amygdala volume and age peaking at ~16 years of age (14), this was not present in the current sample which may be due to the sample being skewed towards younger participants (ages 6-18). Fractional anisotropy and mean diffusivity of the right UF and CB showed a positive linear relationship with age across the sample which is consistent with reports of prolonged maturation of these two white matter tracts (15–18). Lastly, when examining the trajectory of externalizing and internalizing behavior, there was a positive relationship between internalizing behavior and age which was best described by a linear model. This is consistent with reports suggesting that internalizing behavior increases throughout childhood and adolescence peaking at young adulthood (19). There was also a negative linear relationship between externalizing behavior and age which is consistent with previous studies (20). Overall, we find consistent relationships of the trajectories of the brain and behavioral measures of interest across age between the current sample and prior reports. This provides confidence that the current sample was suitable for detecting previously reported brain-behavior relationships. As a result, the null results of this study are unlikely the result of a dataset incompatible with prior reports.

### 7. Post-hoc Bootstrap Resampling Analysis

Stability of the parameter estimates of the null brain-behavior models were assessed using a case-resampling bootstrap (Monte Carlo) approach. Using this approach, 1000 iterations of each data matrix were generated and used to perform the linear regression models and extract three model parameters of interest: regression coefficients, t-statistic, and Cohen's D. The standard error of the regression coefficient was used to assess model stability and the distribution of the three model parameters were used to assess reliability of the models.

#### *7.1. Structural Covariance and Functional Connectivity Bootstrap Resampling*

For the structural covariance and functional connectivity models, RStudio was used to generate 1000 design matrices with replacement. Each design matrix was submitted to a separate linear regression analysis via PALM using one permutation and no TFCE to extract the regression coefficient, t-statistic, and Cohen's D effect size at each vertex. Once all 1000 models were performed in PALM, a matrix was generated in which the rows encompassed the model parameter of interest (regression coefficient, t-statistic or Cohen's D) at each vertex (>50,000 rows) and the columns were the bootstrapped resampled parameters from each model (1000 columns). RStudio was then used to calculate the bootstrap resampled standard error of the regression coefficient at each vertex based on the 1000 iterations. In addition to the standard errors, the mean regression coefficient, t-statistic and effect size (averaged across all the bootstrapped resamples) were calculated. See Figure S9 and S10 for more results from this analysis.

#### *7.2. White Matter Connectivity Bootstrapping*

Bootstrap resampling for the white matter connectivity models were carried out in RStudio using the *Boot* function in the *car* package. This function generated 1000 data matrices and performed the baseline linear regression model on each of the generated matrices. Standard errors of the regression coefficients were extracted from each model (*Table S5*-6).

### 8. Tables

*Table S1. Demographic Characteristics of the analyzed rs-fMRI and DWI data*

| Characteristic | Total | ASD | ADHD | OCD | TDC | X2 | p | X |
|---|---|---|---|---|---|---|---|---|
| **rs-fMRI SAMPLE** | | | | | | | | |
| N | 299 | 113 | 85 | 50 | 51 | | | |
| Males | 214 | 88 | 61 | 31 | 34 | 4.04 | 0.26 | |
| | Mean(SD) | Mean(SD) | Mean(SD) | Mean(SD) | Mean(SD) | F | p | post-hoc tests |
| Age (in years) | 11.83(2.87) | 12.13(3.16) | 11.1(2.48) | 12.9(2.41) | 11.33(2.91) | 5.27 | 0.0014 | ADHD,TDC<ASD,OCD |
| CBCL Total T Score | 61.18(12.14) | 65.85(8.29) | 65.43(9.72) | 61.68(9.15) | 43.92(9.41) | 43.14 | <0.001 | TDC<ASD, ADHD, OCD |
| CBCL Externalizing Behavior T Score | 55.96(12.54) | 59.89(11.01) | 60.85(11.36) | 52.68(11.08) | 42.7(7.5) | 22.54 | <0.001 | TDC<OCD<ASD,ADHD |
| CBCL Internalizing Behavior T Score | 61.35(11.11) | 64.49(8.57) | 62.73(10.32) | 65.59(9.64) | 48.52(9.31) | 23.26 | <0.001 | TDC<ASD, ADHD, OCD |
| Full Scale IQ (age-dependent) | 101.32(19.31) | 93.95(21.9) | 102.3(13.4) | 112.6(20.9) | 111.2(10.4) | 14.19 | <0.001 | ASD<ADHD<OCD, TDC |
| **DWI SAMPLE** | | | | | | | | |
| N | 157 | 78 | 38 | 31 | 10 | | | |
| Males | 119 | 62 | 32 | 22 | 3 | 13.87 | 0.003 | |
| | Mean(SD) | Mean(SD) | Mean(SD) | Mean(SD) | Mean(SD) | F | p | post-hoc tests |
| Age (in years) | 11.48(2.8) | 11.89(3.03) | 10.13(1.95) | 12.31(2.32) | 10.89(3.48) | 4.9 | 0.003 | ADHD<TDC<ASD<OCD |
| CBCL Total T Score | 63.18(11.34) | 65.17(9.18) | 66.5(9.13) | 62.13(9.16) | 38.3(10.5) | 10.63 | <0.001 | TDC<ASD, ADHD, OCD |
| CBCL Externalizing Behavior T Score | 57.47(11.62) | 58.65(11.03) | 61.74(9.16) | 54.77(11.36) | 40.4(9.33) | 5.18 | 0.002 | TDC<ASD, ADHD, OCD |
| CBCL Internalizing Behavior T Score | 62.38(11.08) | 64.15(10.43) | 61.34(9.5) | 65.29(9.79) | 43.5(7.04) | 7.48 | <0.001 | TDC<ASD, ADHD, OCD |
| Full Scale IQ (age-dependent) | 98.9(19.15) | 95.47(20.1) | 99.58(14.59) | 113.87(24.6) | 110(8.95) | 3.79 | 0.01 | ASD,ADHD<OCD, TDC |

*Table S2. Relationship between brain and behavior metrics and age*

| Measure of Interest | Intercept | Linear | Quadratic | Adjusted.R.squared | P.value | DoF |
|---|---|---|---|---|---|---|
| **Structural Index** | | | | | | |
| Left Amygdala Volume | 30.190 | F=26.75 | | 0.070 | <0.001 | 1,323 |
| Right Amygdala Volume | 31.140 | F=21.06 | | 0.060 | <0.001 | 1,323 |
| Left Cortical Thickness | 95.950 | F= 41.8 | | 0.112 | <0.001 | 1,323 |
| Right Cortical Thickness | 94.190 | F=44.05 | | 0.117 | <0.001 | 1,323 |
| **Diffusion Index** | | | | | | |
| FA left Cingulum Bundle | 31.400 | F=33.6 | | 0.172 | <0.001 | 1,155 |
| FA right Cingulum Bundle | 31.450 | F=33.5 | | 0.170 | <0.001 | 1,155 |
| MD left Cingulum Bundle | 95.750 | F=57.01 | | 0.264 | <0.001 | 1,155 |
| MD right Cingulum Bundle | 95.700 | F=57 | | 0.300 | <0.001 | 1,555 |
| FA left Uncinate Fasciculus | 6.370 | | F=1.15 | 0.002 | 0.32 | 2,154 |
| FA right Uncinate Fasciculus | 35.262 | F=0.162 | | -0.005 | 0.69 | 1,155 |
| MD right Uncinate Fasciculus | 63.900 | F=17.04 | | 0.090 | <0.001 | 1,155 |
| MD left Uncinate Fasciculus | 58.007 | F=7.637 | | 0.041 | 0.006 | 1,155 |
| **Behavioral Index** | | | | | | |
| Externalizing Behavior | 6.860 | F=4.03 | | 0.009 | 0.04 | 1,323 |
| Internalizing Behavior | 2.970 | F=11.3 | | 0.031 | <0.001 | 1,323 |

*Table S3. Previous studies examining the relationship between the cortico-amygdalar network and externalizing and internalizing behaviour in developmental cohorts.*

| Author | Year Published | Sample Population | Age Range | Sample Size | Behavioural Measure | Score Range | Brain Metric | Main Finding | Measure of Effect |
|---|---|---|---|---|---|---|---|---|---|
| Saxbe et al | 2018 | TDC | 11.79 – 13.93 | 21 | self-report YSR and CBCL | 33– 73 (T-score) | cortico-amygdalar resting state functional connectivity | Externalizing behaviour linked to stronger connectivity between the amygdala and the sgACC, OFC, insula And mPFC in mid-adolescence | unable to extract effect size |
| Ali et al | 2019 | TDC | 7 years old | 47 | Internalizing Behaviour from CBCL | Internalizing Raw Scores: 3 years = 3.5±0.3 6 years = 6 ±0.31 Eight years = 8.7 ±0.71 | FA of uncinate fasciulus and cingulum bundle | FA of cingulum bundle significantly associated with internalizing symptoms. Radial diffusivity of right uncinate fasciculus positvely linked to internalizing behaviour | Cingulum: r-=0.35, r=-0.338 Uncinate Fasciculus: r=0.4 |
| Andre et al | 2019 | TDC | 6 – 16 | 48 | BASC-2 | Internalizing behaviour: 154 ± 26 Externalizing behaviour: 157 ± 25 | Gray matter volume and diffusion metrics of the cortico-limbic regions | Externalizing behavior linked to FA of the left cingulum And externalizing behaviour. Age-FA of uncinate fasiculus interaction associated with externalizing scores | left cingulum = d=0.79 Age-agression on left UF = d=0.734 |
| Karlsgodt et al | 2018 | TDC | 10 – 19 | 53 | self-report YSR and CBCL | Internalizing T-score : 27-69 Externalizing T-score: 27-72 | MSIT Cognitive Control Task functional connectivity | Internalizing and externalizing behavior associated with hyperactivity of mPFC and parietal lobe, respectively | unable to extract effect size |
| Ibrahim et al | 2019 | ASD and TDC | 8 – 16 | 57 (ASD + disruptive behaviour, n=18: ASD, n =20: TDC, n=19) | Externalizing behaviour from CBCL | Externalizing behaviour (T-score): ASD+BD = 69.3±5.8 ASD = 44.9 ±7.1 TD = 41.1±7.8 Internalizing behaviour (T-score): ASD+DB = 67.1±8.6 ASD = 54.3±6.9 TD = 43.1±6.5 | Cortico-amygdalar task based functional connectivity | Externalizing behaviour associated with lower amygdala-dlPFC functional connectivity in patients with ASD. Positive relationship between externalizing scores and amygdala response in ASD+DB group | Amygdala-vlPFC: semipartial r = -0.397 Amygdala: semipartial r = 0.318 |
| Chabernaud et al | 2012 | ADHD and TDC | 7.2 – 13.3 | 37 TDC, 37 ADHD | Internalizing and Externalizing Scores from CBCL | Externalizing T-score: 33-78 Internalizing T-score: 34-74 | resting state functional connectivity: DMN ROIs | Internalizing and externalizing behavior was linked to alterations of the default mode network across children with ADHD and TDC | unable to extract effect size |
| Qin et al | 2014 | TDC | 7 – 9 | 76 | Anxiety/depressed subscale from CBCL | 50-72 (anxiety subscale T-score) | Amygdala volume and cortico-amygdalar resting state functional connectivity | Children with greater anxiety had larger amygdala, vmPFC, and insula volume | Amygdala: r=0.42, r=0.23 VmPFC: r=0.46 Insula: 0.54 |
| van der Plas et al | 2010 | TDC | 7 – 17 | 116 | Fearfulness subscale of the Pediatric Behavior Scale | Boys = 1.62 ± 1.52 Girls = 1.71 ± 1.23 | Amygdala volume | significant positive correlation between Right amygdala volume and fearfulness scores in girls | r=0.29 |
| Vijayakumar et al | 2016 | TDC | 11 – 20 | 166 | CESD | CESD = 31.33±9.61 BAI = 8.36±8.52 | cortico-amygdalar structural covariance | Significant interaction between age-right amygdala Volume change predicting left anterior cortical thickness in relation to CESD scores | Cohen's d=0.112 |
| Albaugh et al | 2016 | TDC | 5.7 – 18.4 | 175 | Anxiety/depression subscale of CBCL | 1.66 ±1.89 (raw score) | Fractional anisotropy of white matter | age-by-CBCL anxious/depression subscale linked to FA of the left and right cingulum | Left cingulum bundle: cohens d = 0.18 Right cingulum bundle: cohens d = 0.23 |
| Ducharme et al | 2011 | TDC | 6 – 18 | 193 | Aggressive Behaviour Subscale of CBCL | 2.47±0.19 (aggression subscale raw score) | Cortical Thickness and subcortical volumes | Aggression CBCL scores associated with a thinner ACC cortex | unable to extract effect size |
| Snyder et al | 2017 | TDC | 6 – 10 | 254 | CBCL, CBQ | Internalizing raw score: 2 64 ± 2.07 Externalizing raw score: 3.58 ± 2.57 | Gray matter volume | Higher levels of internalizating factors associated with smaller amygdala and insula volumes | Amygdala: B= -0.25 Insula: B = -0.31, -0.36 |
| Ameis et al | 2014 | TDC | 6 – 18 | 291 | Parent-reported Externalizing CBCL Scores | 32-69 (T-scores) | cortico-amygdalar structural covariance | cortico-amygdalar covariance relationship with externalizing behavior | Cohen's d=0.244 |

*Note:*
Some studies completed their analyses using softwares (e.g. FSL, SPM) and as a result did not report specific statistical metrics. For those studies, we were unable to calculate effect size.
TDC = typically developing children, ASD = Autism Spectrum Disorder, ADHD = Attention-Deficit/Hyperactivity Disorder, YSR = Youth Self Report, CBCL = Child Behavior Checklist, CESD = Centre for Epidemiological Studies - Depression,
CBQ = Children's Behavior Quesionnaire, sgACC = subgenual anterior cingulate cortex, OFC = orbitofrontal cortex, mPRC = medial prefrontal cortex, dlPFC = dorsolateral prefrontal cortex, vmPFC = ventromedial prefrontal cortex,
vlPFC = ventrolateral prefrontal cortex, FA = fractional anisotropy

*Table S4. Subset of the T1w sample where the presence of confirmed co-occurring diagnoses was available*

| **Co-occurring Diagnoses** | **Diagnosis** | | |
|---|---|---|---|
| | **ASD** | **ADHD** | **OCD** |
| ASD | - | 1 | 0 |
| ADHD | 6 | - | 2 |
| OCD | 0 | 1 | - |
| Anxiety Disorders | 3 | 9 | 8 |
| Tic Disorder | 1 | 2 | 2 |
| Disruptive Behavior Disorders (Conduct Disorder, Oppositional Defiance Disorder, Other) | 0 | 16 | 0 |
| Intellectual Disability | 1 | 0 | 0 |
| IQ < 70 | 15 | 0 | 1 |

Note: ASD = Autism Spectrum Disorder, ADHD = Attention-Deficit/Hyperactivity Disorder, OCD = Obsessive Compulsive Disorder. Comorbidity data was collected using a clinician completed checklist. Confirmed diagnoses are included in this table.

Table S5. Results of the case-resampling bootstrap analysis of the white matter connectivity linear regression models (main sample; n=157).

| **Brain Metric** | **Behavioral Metric** | **Bootstrap 95% CI (X, X)** | **Bootstrap Standard Error** |
|---|---|---|---|
| Left CB FA | Externalizing Behavior | (-0.00049, 0.00046) | 0.00024519 |
| Left CB MD | Externalizing Behavior | (-9.79e-08, 7.66e-07) | 2.20e-07 |
| Left CB FA | Internalizing Behavior | (-0.00071, 0.00039) | 0.00028315 |
| Left CB MD | Internalizing Behavior | (-2.53e-07, 5.14e-07) | 1.96e-07 |
| Right CB FA | Externalizing Behavior | (-0.00049, 0.00048) | 0.00024968 |
| Right CB MD | Externalizing Behavior | (-1.086e-07, 7.52e-07) | 2.20e-07 |
| Right CB FA | Internalizing Behavior | (-0.0007, 0.0004) | 0.00027334 |
| Right CB MD | Internalizing Behavior | (-3.12e-07, 4.68e-07) | 1.99e-07 |
| Left UF FA | Externalizing Behavior | (-0.0006, 0.0006) | 0.00029866 |
| Left UF MD | Externalizing Behavior | (-4.74e-07, 8.17e-07) | 3.29e-07 |
| Left UF FA | Internalizing Behavior | (-0.0003, -0.0003) | 0.00034875 |
| Left UF MD | Internalizing Behavior | (-8.97e-07, 4.87e-07) | 3.53E-07 |
| Right UF FA | Externalizing Behavior | (-0.00053, 0.00058) | 0.00028703 |
| Right UF MD | Externalizing Behavior | (-2.83e-07, 1.40e-06) | 4.31E-07 |
| Right UF FA | Internalizing Behavior | (-0.0006, 0.0006) | 0.00032562 |
| Right UF MD | Internalizing Behavior | (-4.556992e-07, 6.799967e-07) | 2.90E-07 |

Table S6. Results of the case-resampling bootstrap analysis of the sensitivity white matter connectivity linear regression models (subsample; n=110).

| Brain Metric | Behavioral Metric | Bootstrap 95% CI (X, X) | Bootstrap Standard Error |
|---|---|---|---|
| Left CB FA | Externalizing Behavior | (-0.0006, 0.00059) | 0.00032634 |
| Left CB MD | Externalizing Behavior | (-4.5e-08, 1.11e-06) | 2.94E-07 |
| Left CB FA | Internalizing Behavior | (-4.48e-08, 1.11e-06) | 0.00031241 |
| Left CB MD | Internalizing Behavior | (-2.18e-09, 7.78e-07) | 1.99E-07 |
| Right CB FA | Externalizing Behavior | (-0.0007, 0.0006) | 0.00033458 |
| Right CB MD | Externalizing Behavior | (-7.22e-08, 1.11e-06) | 3.02E-07 |
| Right CB FA | Internalizing Behavior | (-0.0011, 0.00011) | 0.00032717 |
| Right CB MD | Internalizing Behavior | (2.06e-09, 8.06e-07) | 2.05E-07 |
| Left UF FA | Externalizing Behavior | (-0.0007, 0.0006) | 0.0003491 |
| Left UF MD | Externalizing Behavior | (-5.75e-07, 1.29e-06) | 4.77E-07 |
| Left UF FA | Internalizing Behavior | (-0.0003, 0.0011) | 0.00037528 |
| Left UF MD | Internalizing Behavior | (-8.44e-07, 8.49e-07) | 4.32E-07 |
| Right UF FA | Externalizing Behavior | (-0.00099, 0.00055) | 0.00039488 |
| Right UF MD | Externalizing Behavior | (-6.15e-08, 2.07e-06) | 5.43E-07 |
| Right UF FA | Internalizing Behavior | (-0.0008, 0.00066) | 0.00037251 |
| Right UF MD | Internalizing Behavior | (2.447816e-08, 1.016086e-06) | 2.53E-07 |

**9. Figures**

*Figure S1. The a priori visual QC rating system structure used to assess and rate the quality of T1-weighted brain images. Participants were excluded if they received a score of 4 or 5.*

### ARTIFACT RATING SYSTEM STRUCTURE

| 1 = NO/MILD | 2 = MILD/MODERATE | 3 = MODERATE | 4 = MODERATE/SEVERE | 5 = SEVERE |
|---|---|---|---|---|
| • No visible artefacts<br>• Very light rings in certain areas | • Light rings in more broad regions of the brain (e.g. across cortex) | • Rings in more broad regions of the brain (e.g. across cortex)<br>• Minor segmentation errors with pial and white matter layers | • Relatively prominent and dark rings in most of the brain regions<br>• Segmentation errors with pial and white matter layers<br>• Minor resulting distortions<br>• Minor ghosting<br>• Localized white areas (indicative of intensity non-uniformity) in the brain scan (such as small areas of the cortex) | • Prominent and dark rings in most of the brain regions<br>• Segmentation errors with pial and white matter layers<br>• Extreme distortions (such as missing brain regions, white sections in brain scan etc.)<br>• Full ghosting |

*Figure S2. Diagrams presenting the subsample which includes children with autism spectrum disorder (ASD), attention-deficit/hyperactivity disorder (ADHD), obsessive compulsive disorder (OCD) and typically developing children (TDC) scanned at the Hospital for Sick Children as of January 2020. Imaging data from T1-weighted (T1w), resting state fMRI (rsfMRI) and diffusion weighted imaging (DWI) sequences are presented. The reasons for exclusion presented for level 1: participants being outside the 6-18 age range at time of scan, a greater than 1 month time gap between scan and CBCL administration, and missing CBCL data; level 2: persistent processing errors at any point within the processing pipeline (e.g. errors in the fMRIprep pipeline); level 3: exclusion based on quality control (QC; details presented in the paper and supplement). The numbers for the final analysed sample for each imaging modality are presented. For the T1w and rs-fMRI samples, participants were scanned on a 3T Siemens Tim Trio scanner prior to June 2016 when the scanner was upgraded to the PrismaFIT. For rs-fMRI acquisitions, participants scanned on the Tim Trio selected a movie to watch and participants scanned on the PrismaFIT viewed a naturalistic film (inscapes). The study includes only single-shell DWI acquisitions (n=262) completed on the Tim Trio scanner.*

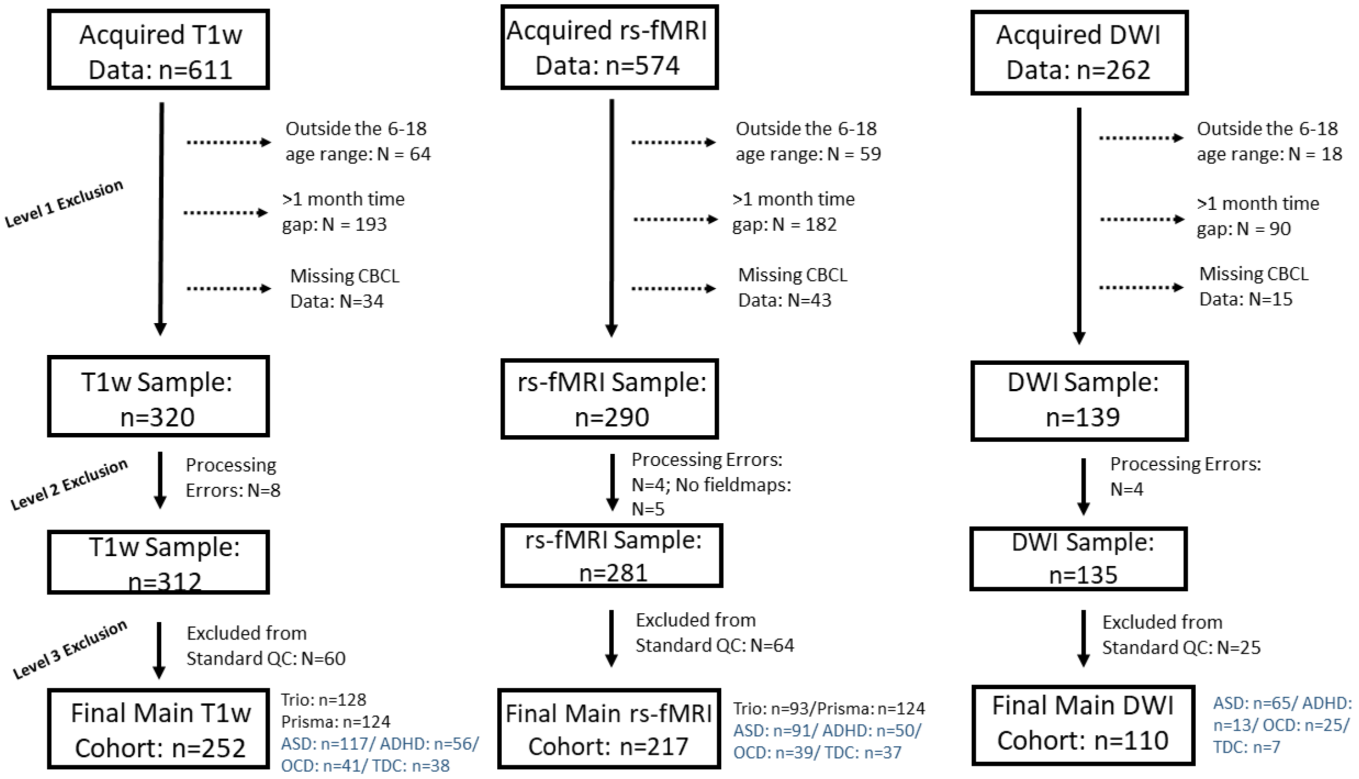

*Figure S3. Unthresholded spatial p-map output of the relationship between externalizing/internalizing behaviors and cortico-amygalar structural covariance and functional connectivity in the planned subsample analysis. A logp value of 1.6 is considered significant and none of the results reached that value.*

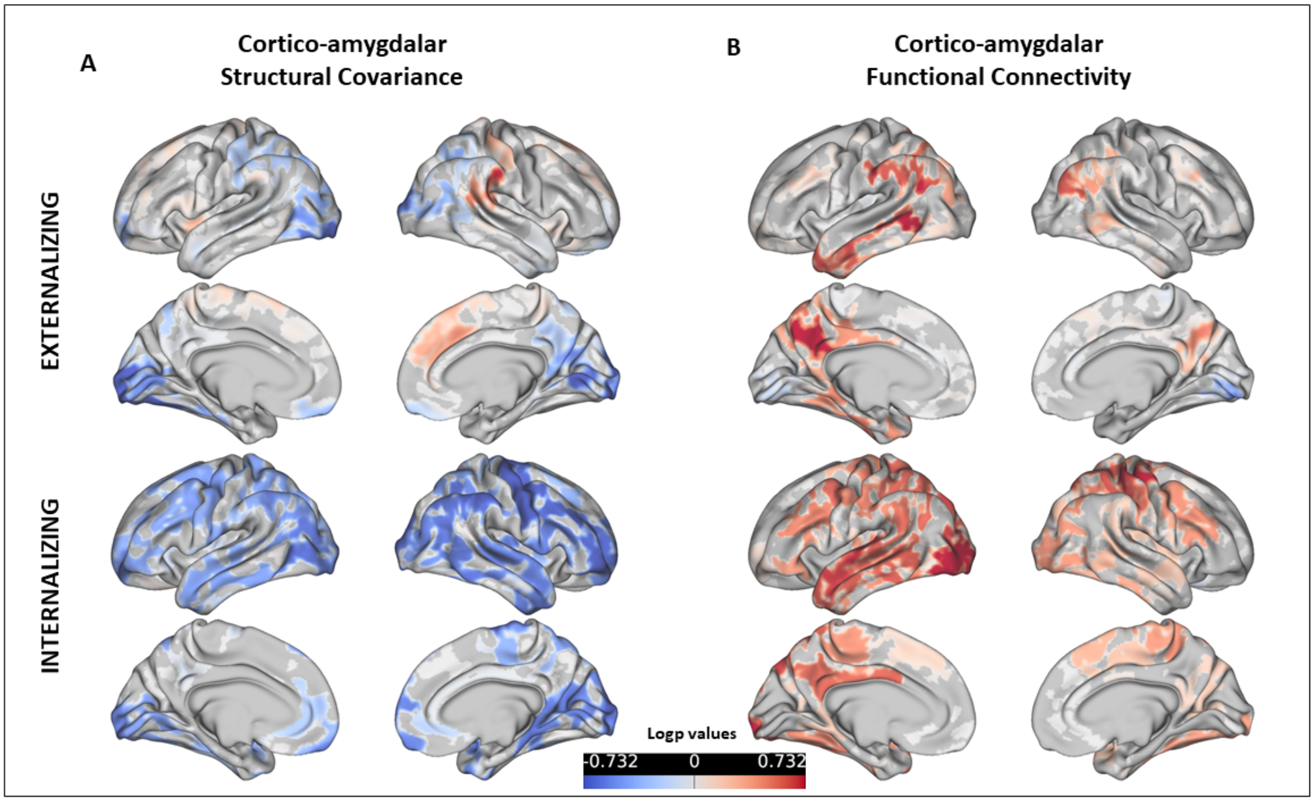

*Figure S4. Unthresholded p-map depicting relationship between cortico-amygdalar covariance (right amygdala volume*right superior temporal gyrus thickness) and externalizing behavior across the sample (n=346). This region is depicted as it showed a stronger brain-behavior relationship (relative to other regions, but it remained non-significant). High and low externalizing groups were determined using a median split (median = 10). These results did not meet the significance threshold.*

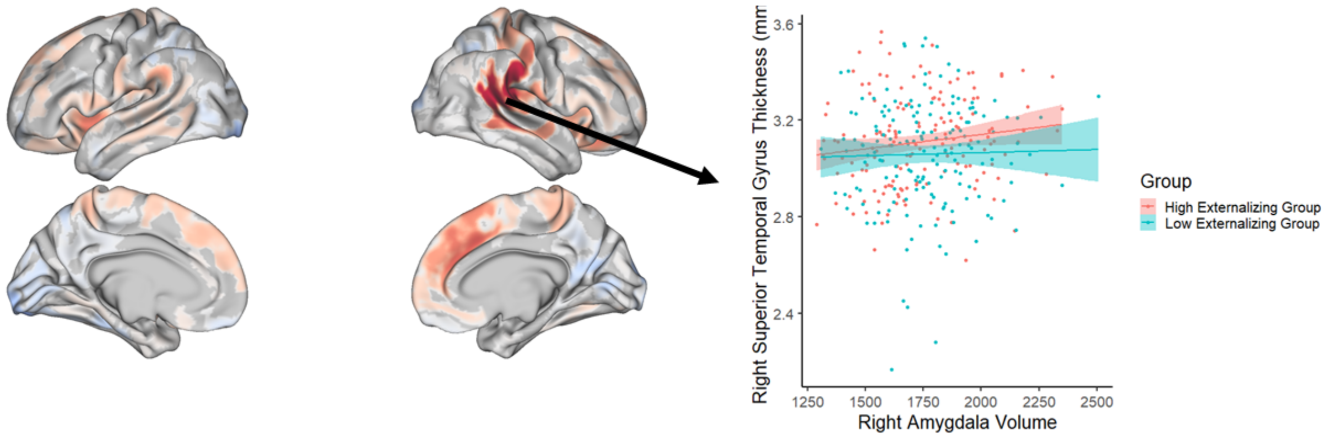

*Figure S5. Unthresholded p-map depicting relationship between cortico-amygdalar covariance (left amygdala volume*left middle frontal gyrus thickness) and internalizing behavior across the sample (n=346). This region is depicted as it showed a stronger relationship (relative to other regions, but it remained non-significant). High and low internalizing groups were determined using a median split (median = 12). These results did not meet the significance threshold.*

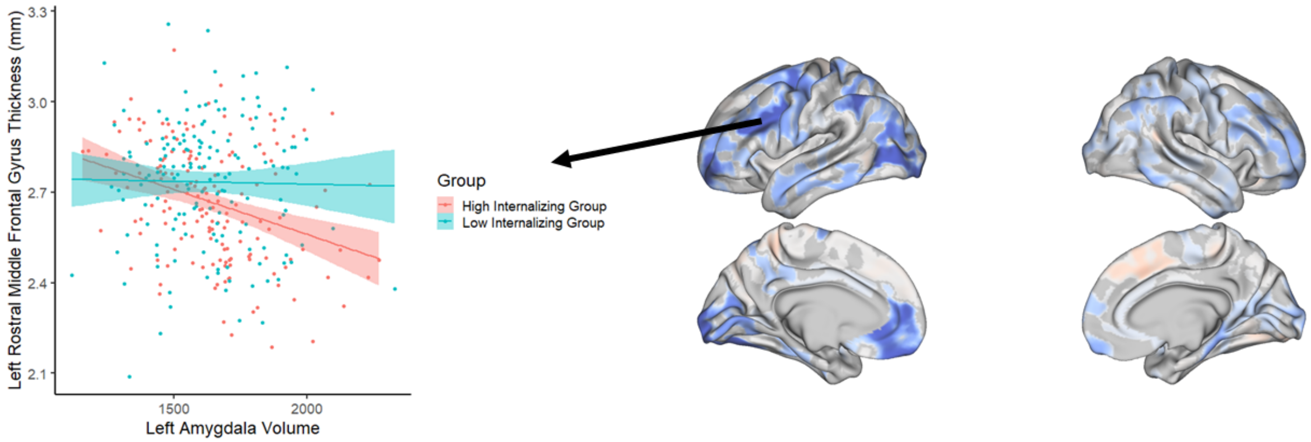

*Figure S6a. Right hemisphere cortical thickness and amygdala volume presented across the age range of the analysed sample.*

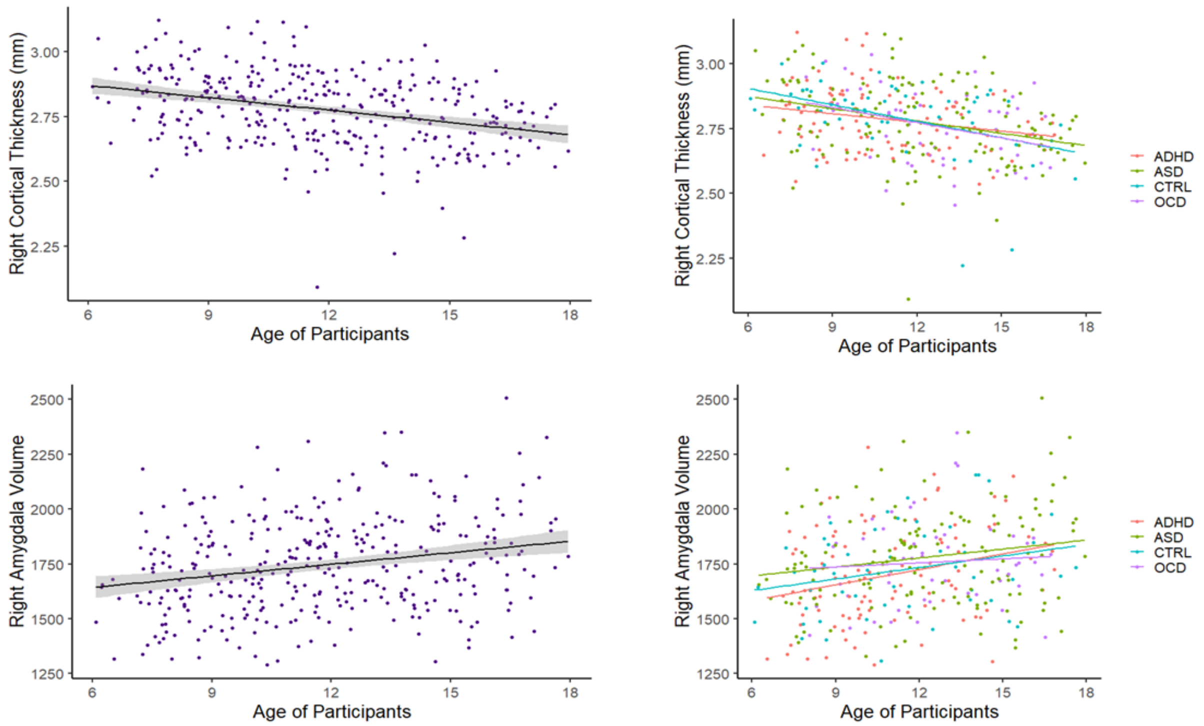


Note: Cortical thickness and amygdala volume were parcellated from the Desikan-Killiany Atlas. Right cortical thickness is an average of the thickness of all the cortical regions-of-interest of the right hemisphere. ADHD = Attention-Deficit/Hyperactivity Disorder, ASD = Autism Spectrum Disorder, CTRL = healthy control (i.e. typically developing), OCD = Obsessive Compulsive Disorder

*Figure S6b. FA and MD (units: mm2/s) of the right uncinate fasciculus presented across the age range of the analysed sample.*

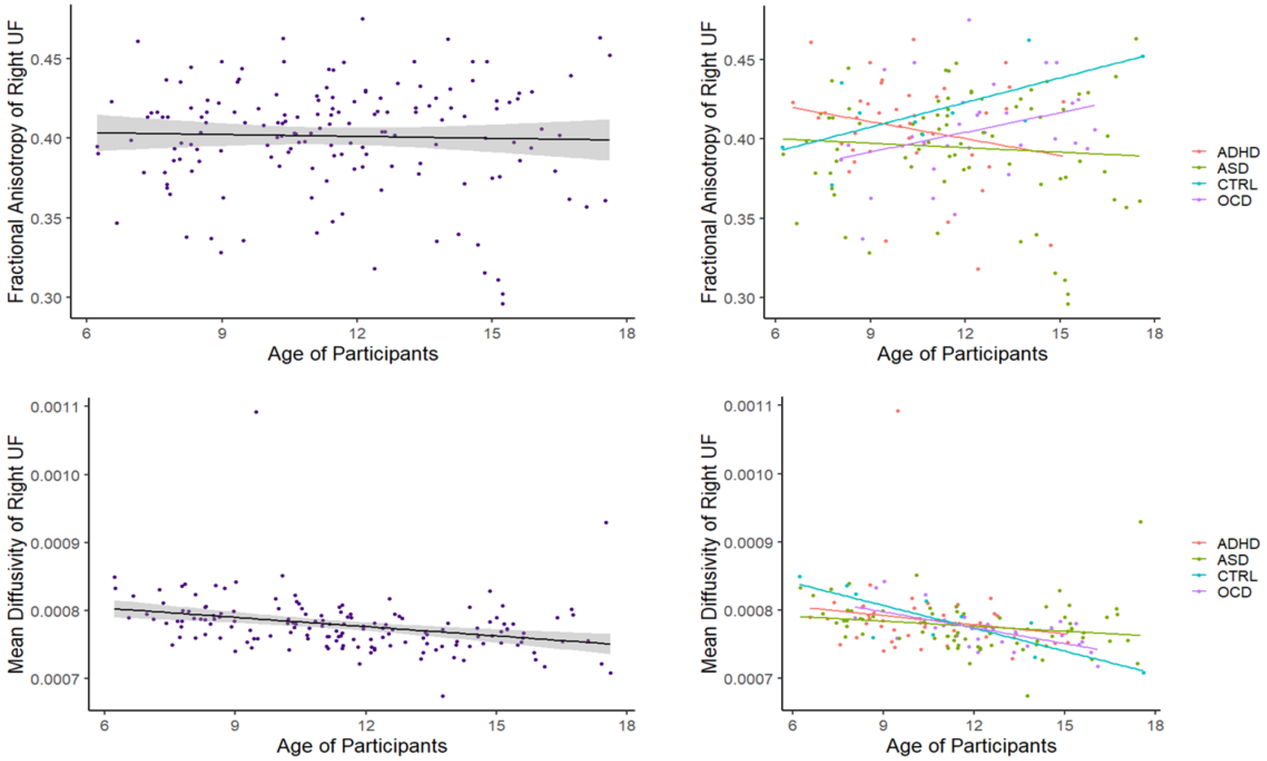


Note: UF = Uncinate Fasciculus; ADHD = Attention-Deficit/Hyperactivity Disorder, ASD = Autism Spectrum Disorder, CTRL = healthy control (i.e. typically developing), OCD = Obsessive Compulsive Disorder

*Figure S6c. FA and MD (units: mm2/s) of the right cingulum bundle presented across the age range of the analysed sample.*

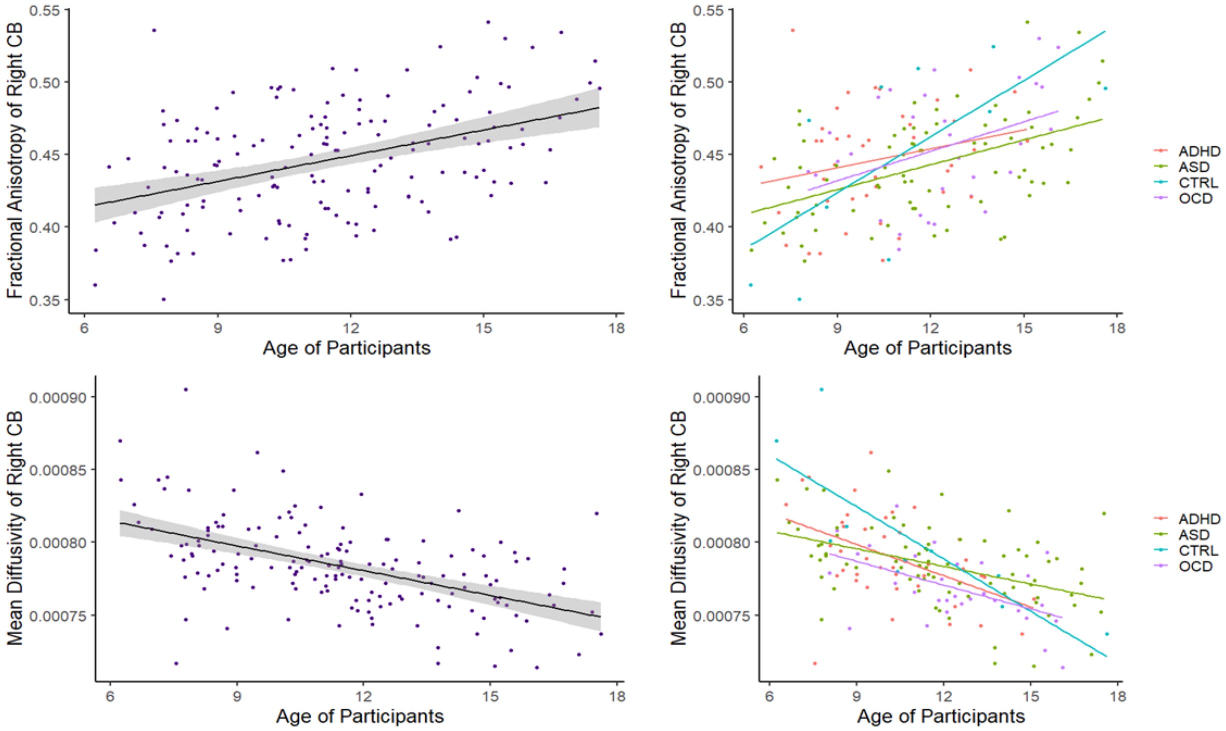


Note: CB = Cingulum Bundle; ADHD = Attention-Deficit/Hyperactivity Disorder, ASD = Autism Spectrum Disorder, CTRL = healthy control (i.e. typically developing), OCD = Obsessive Compulsive Disorder

*Figure S7. Relationship between mean diffusivity (units: mm2/s) of the right uncinate fasciculus and externalizing behavior before ($F=8.96$, $p_{Model} <0.001$, $t_{Externalizing} = 3.43$, $p_{Externalizing} = 0.009$) and after ($F=9.4$, $p_{Model} <0.001$, $t_{Externalizing} = 1.19$, $p_{Externalizing} = 0.24$) outlier removal in the DWI subsample (n=110). Outliers were removed based on whether their mean diffusivity was two standard deviations from the sample mean.*

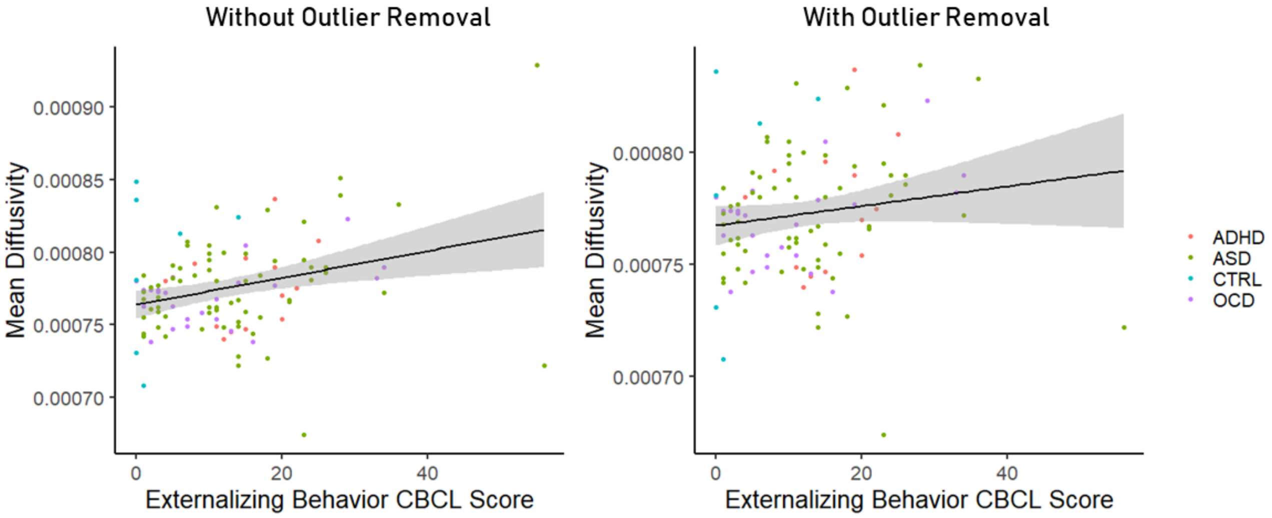

*Figure S8. Relationship between mean diffusivity (units: mm2/s) of the right cingulum bundle and externalizing behavior in the DWI subsample (n=110) prior to multiple comparison correction.*

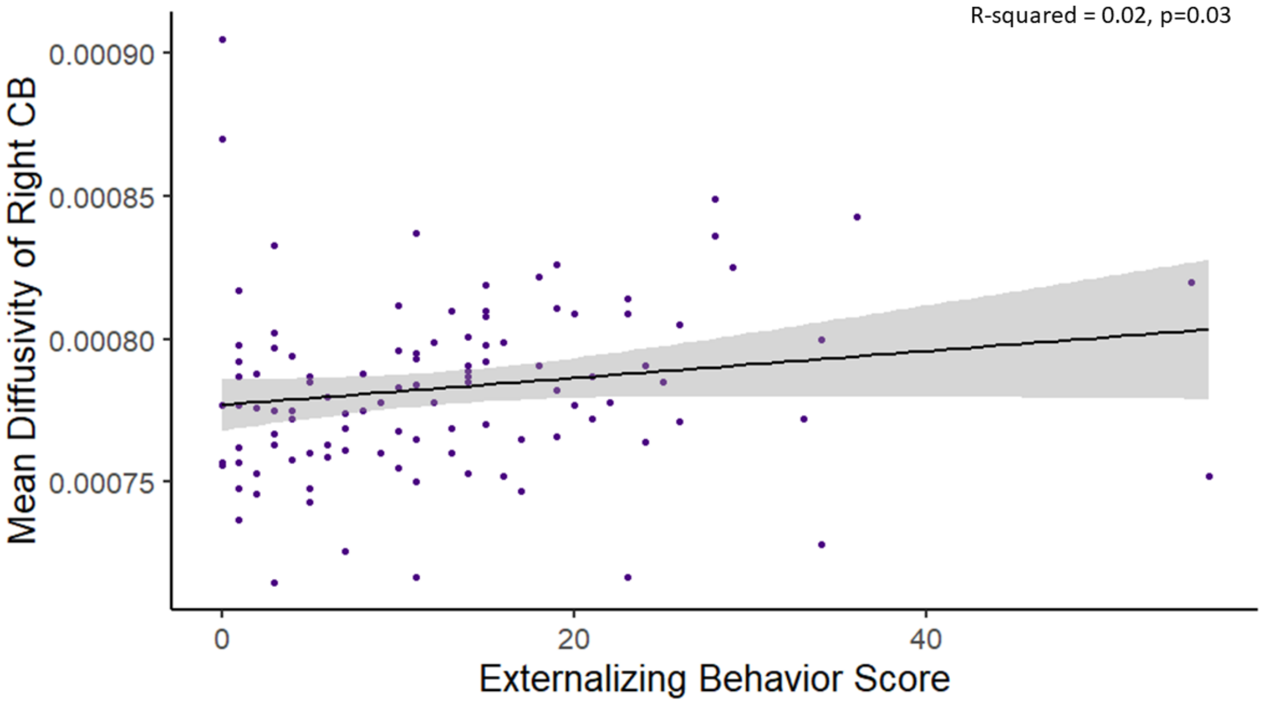

*Figure S9. Bootstrap resampling of the remaining structural covariance and functional connectivity models. Scatterplots between the bootstrapped mean regression coefficients (averaged across 1000 resamples x-axis) and bootstrapped standard error (y-axis) are shown. Each point on the figure is one vertex. Points within the brown circle indicate the vertices with a t-statistic greater than 4 to illustrate a proxy for 'higher signal' and low noise vertices. Points not within the brown circle depict the vertices with a t-statistic less than 4 to illustrate 'low signal' and low noise vertices. The standard errors of the regression coefficients are near zero for all vertices across structural covariance and functional connectivity models.*

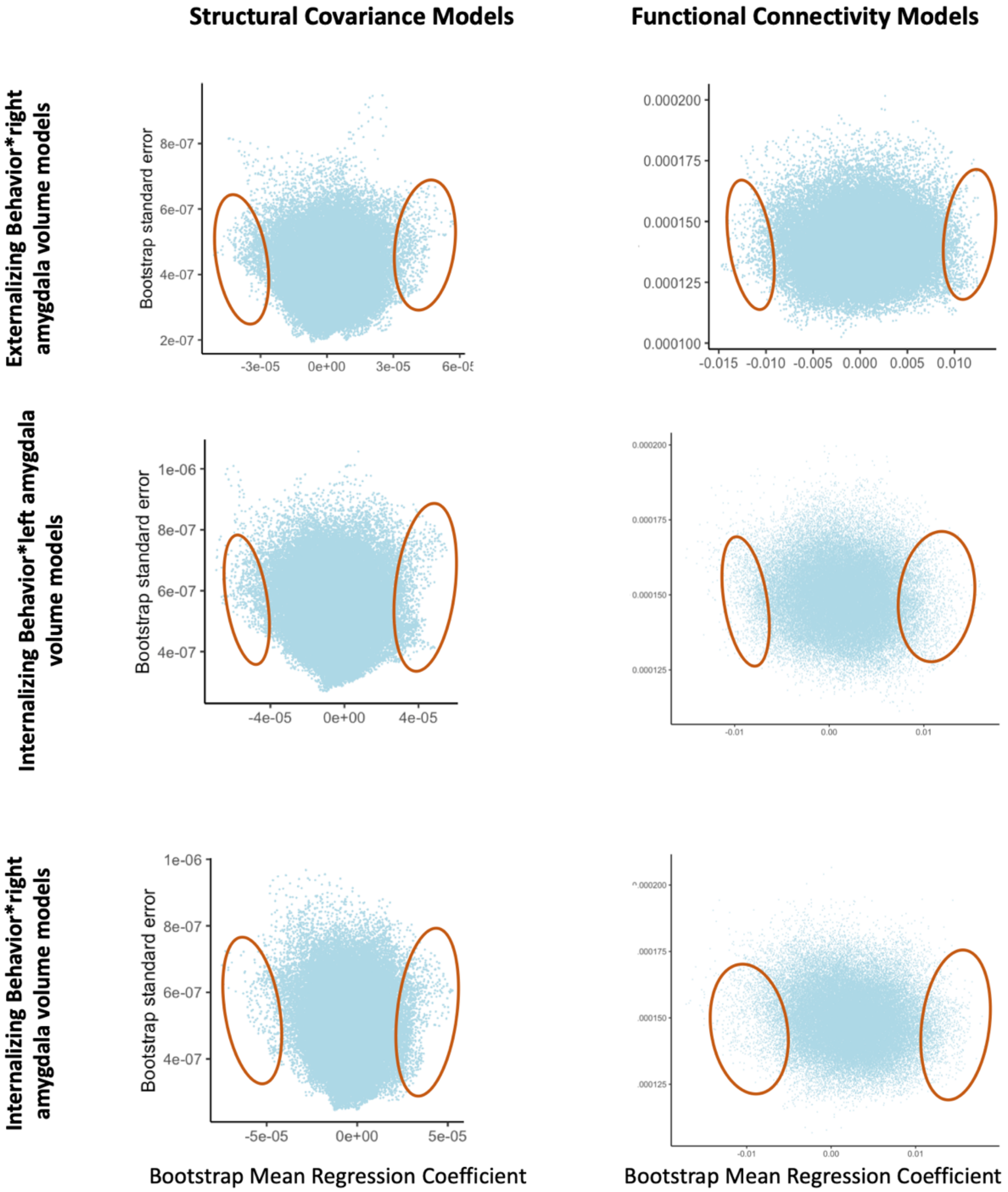

*Figure S10. Bootstrap resampling of the internalizing behavior*right amygdala structural covariance and functional connectivity models among the subsample (n=252, n=217). This figure depicts the results of a bootstrap resampling analysis in which 1000 iterations of each data matrix were generated and used to perform the linear regression models in PALM. Panel A illustrates scatterplots between the mean regression coefficient (averaged across 1000 resamples) and the bootstrapped standard errors of the regression coefficients of each vertex for the structural covariance and functional connectivity models. Points colored in pink depict the vertices with a t-statistic greater than 4 to illustrate a proxy for 'higher signal' and low noise vertices. Points colored in blue depict the vertices with a t-statistic less than 4 to illustrate 'low signal' and low noise vertices. Panel B depicts the histogram of the mean regression coefficients of each vertex across the 1000 bootstrapped resampled analyses. Panel C depicts the histogram of the mean t-statistic of each vertex across the 1000 bootstrapped resampled analyses. Panel D depicts the histogram of the mean effect size of the model at each vertex across the 1000 bootstrapped resampled analyses.*

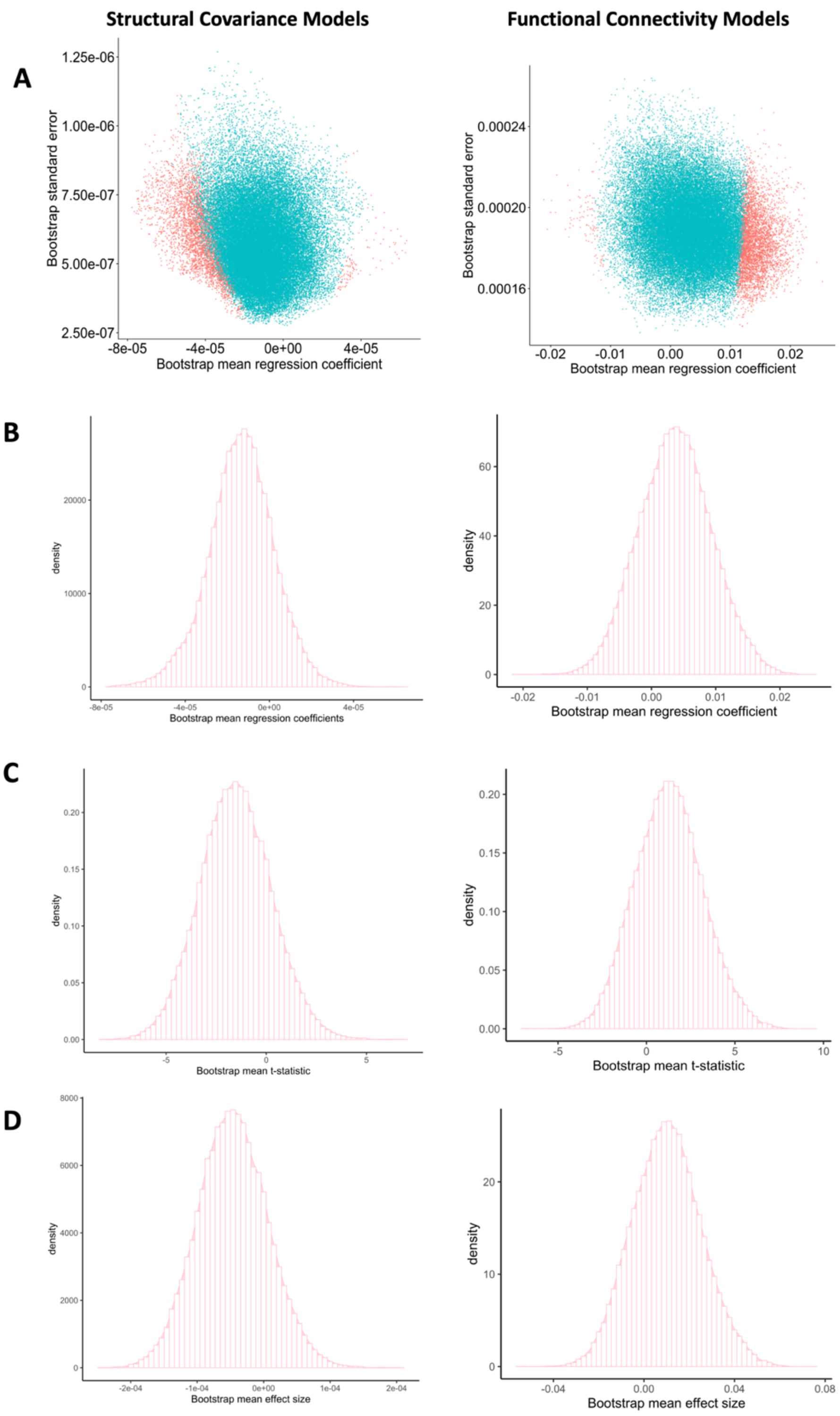
Structural Covariance Models
Functional Connectivity Models
A
Bootstrap standard error
Bootstrap mean regression coefficient
1.25e-06
1.00e-06
7.50e-07
5.00e-07
2.50e-07
-8e-05
-4e-05
0e+00
4e-05
0.00024
0.00020
0.00016
-0.02
-0.01
0.00
0.01
0.02
B
density
Bootstrap mean regression coefficients
Bootstrap mean regression coefficient
C
density
Bootstrap mean t-statistic
D
density
Bootstrap mean effect size

https://doi.org/10.5281/zenodo.3700055.
3. Gorgolewski K, Burns CD, Madison C, Clark D, Halchenko YO, Waskom ML, Ghosh SS. Nipype: a flexible, lightweight and extensible neuroimaging data processing framework in python. Front Neuroinform. 2011 Aug 22;5(August):13. doi:10.3389/fninf.2011.00013.
4. Gorgolewski KJ, Esteban O, Ellis DG, Notter MP, Ziegler E, Johnson H, Hamalainen C, Yvernault B, Burns C, Manhães-Savio A, Jarecka D, Markiewicz CJ, Salo T, Clark D, Waskom M, Wong J, Modat M, Dewey BE, Clark MG, Dayan M, Loney F, Madison C, Gramfort A, Keshavan A, Berleant S, Pinsard B, Goncalves M, Clark D, Cipollini B, Varoquaux G, Wassermann D, Rokem A, Halchenko YO, Forbes J, Moloney B, Malone IB, Hanke M, Mordom D, Buchanan C, Pauli WM, Huntenburg JM, Horea C, Schwartz Y, Tungaraza R, Iqbal S, Kleesiek J, Sikka S, Frohlich C, Kent J, Perez-Guevara M, Watanabe A, Welch D, Cumba C, Ginsburg D, Eshaghi A, Kastman E, Bougacha S, Blair R, Acland B, Gillman A, Schaefer A, Nichols BN, Giavasis S, Erickson D, Correa C, Ghayoor A, Küttner R, Haselgrove C, Zhou D, Craddock RC, Haehn D, Lampe L, Millman J, Lai J, Renfro M, Liu S, Stadler J, Glatard T, Kahn AE, Kong X-Z, Triplett W, Park A, McDermottroe C, Hallquist M, Poldrack R, Perkins LN, Noel M, Gerhard S, Salvatore J, Mertz F, Broderick W, Inati S, Hinds O, Brett M, Durnez J, Tambini A, Rothmei S, Andberg SK, Cooper G, Marina A, Mattfeld A, Urchs S, Sharp P,

Matsubara K, Geisler D, Cheung B, Floren A, Nickson T, Pannetier N, Weinstein A, Dubois M, Arias J, Tarbert C, Schlamp K, Jordan K, Liem F, Saase V, Harms R, Khanuja R, Podranski K, Flandin G, Papadopoulos Orfanos D, Schwabacher I, McNamee D, Falkiewicz M, Pellman J, Linkersdörfer J, Varada J, Pérez-García F, Davison A, Shachnev D, Ghosh S. Nipype: a flexible, lightweight and extensible neuroimaging data processing framework in Python. 2017. doi:10.5281/zenodo.581704.
5. Tustison NJ, Avants BB, Cook PA, Zheng Y, Egan A, Yushkevich PA, Gee JC. N4ITK: improved N3 bias correction. IEEE Trans Med Imaging. 2010 Jun;29(6):1310–20. doi:10.1109/TMI.2010.2046908.
6. Dale A, Fischl B, Sereno MI. Cortical Surface-Based Analysis: I. Segmentation and Surface Reconstruction. Neuroimage. 1999;9(2):179–94. doi:10.1006/nimg.1998.0395.
7. Fonov VS, Evans AC, McKinstry RC, Almli CR, Collins DL. Unbiased nonlinear average age-appropriate brain templates from birth to adulthood. NeuroImage; Amsterdam. 2009 Jul 1;47:S102. doi:10.1016/S1053-8119(09)70884-5.
8. Avants BB, Epstein CL, Grossman M, Gee JC. Symmetric diffeomorphic image registration with cross-correlation: evaluating automated labeling of elderly and neurodegenerative brain. Med Image Anal. 2008 Feb;12(1):26–41. doi:10.1016/j.media.2007.06.004.
9. Jenkinson M, Bannister P, Brady M, Smith S. Improved optimization for the robust and accurate linear registration and motion correction of brain images. Neuroimage. 2002 Oct;17(2):825–41. doi:10.1006/nimg.2002.1132.
10. Andersson JLR, Skare S, Ashburner J. How to correct susceptibility distortions in spin-echo echo-planar images: application to diffusion tensor imaging. Neuroimage. 2003 Oct;20(2):870–88. doi:10.1016/S1053-8119(03)00336-7.
11. Cox RW. AFNI: software for analysis and visualization of functional magnetic resonance neuroimages. Comput Biomed Res. 1996 Jun;29(3):162–73. doi:10.1006/cbmr.1996.0014.
12. Jenkinson M. Fast, automated, N-dimensional phase-unwrapping algorithm. Magn Reson Med. 2003 Jan;49(1):193–7. doi:10.1002/mrm.10354.
13. Huntenburg JM. Evaluating nonlinear coregistration of BOLD EPI and T1w images. Freie Universität Berlin; 2014. Available from: http://hdl.handle.net/11858/00-001M-0000-002B-1CB5-A.
14. Wang S, Peterson DJ, Gatenby JC, Li W, Grabowski TJ, Madhyastha TM. Evaluation of Field Map and Nonlinear Registration Methods for Correction of Susceptibility Artifacts in Diffusion MRI. Front Neuroinform. 2017 [cited 2017 Feb 21];11. doi:10.3389/fninf.2017.00017.
15. Treiber JM, White NS, Steed TC, Bartsch H, Holland D, Farid N, McDonald CR, Carter BS, Dale AM, Chen CC. Characterization and Correction of Geometric Distortions in 814 Diffusion Weighted Images. PLoS One. 2016 Mar 30;11(3):e0152472. doi:10.1371/journal.pone.0152472.
16. Greve DN, Fischl B. Accurate and robust brain image alignment using boundary-based registration. Neuroimage. 2009 Oct;48(1):63–72. doi:10.1016/j.neuroimage.2009.06.060.
17. Zhang Y, Brady M, Smith S. Segmentation of brain MR images through a hidden Markov random field model and the expectation-maximization algorithm. IEEE Trans Med Imaging. 2001 Jan;20(1):45–57. doi:10.1109/42.906424.
18. Behzadi Y, Restom K, Liau J, Liu TT. A component based noise correction method (CompCor) for BOLD and perfusion based fMRI. Neuroimage. 2007 Aug 1;37(1):90–101. doi:10.1016/j.neuroimage.2007.04.042.

19. Power JD, Mitra A, Laumann TO, Snyder AZ, Schlaggar BL, Petersen SE. Methods to detect, characterize, and remove motion artifact in resting state fMRI. Neuroimage. 2013 Aug 29;84:320–41. doi:10.1016/j.neuroimage.2013.08.048.
20. Pruim RHR, Mennes M, van Rooij D, Llera A, Buitelaar JK, Beckmann CF. ICA-AROMA: A robust ICA-based strategy for removing motion artifacts from fMRI data. Neuroimage. 2015 May 15;112:267–77. doi:10.1016/j.neuroimage.2015.02.064.
21. Klein A, Ghosh SS, Bao FS, Giard J, Häme Y, Stavsky E, et al. Mindboggling morphometry of human brains. PLoS Comput Biol 13(2): e1005350. 2017. doi:10.1371/journal.pcbi.1005350.
22. Abraham A, Pedregosa F, Eickenberg M, Gervais P, Mueller A, Kossaifi J, Gramfort A, Thirion B, Varoquaux G. Machine learning for neuroimaging with scikit-learn. Front in Neuroinf 8:14. 2014. doi:10.3389/fninf.2014.00014.
